# Taxonomic vs genomic fungi: contrasting evolutionary loss of protistan genomic heritage and emergence of fungal novelties

**DOI:** 10.1101/2022.11.15.516418

**Authors:** Zsolt Merényi, Krisztina Krizsán, Neha Sahu, Xiao-Bin Liu, Balázs Bálint, Jason Stajich, Joseph W. Spatafora, László G. Nagy

## Abstract

Fungi are among the most ecologically important heterotrophs that have radiated into most niches on Earth and fulfil key ecological services. However, despite intense interest in their origins, major genomic trends characterising the evolutionary route from a unicellular opisthokont ancestor to derived multicellular fungi remain poorly known. Here, we reconstructed gene family evolution across 123 genomes of fungi and relatives and show that a dominant trend in early fungal evolution has been the gradual shedding of protist genes and highly episodic innovation via gene duplication. We find that the gene content of early-diverging fungi is protist-like in many respects, owing to the conservation of protist genes in early fungi. While gene loss has been constant and gradual during early fungal evolution, our reconstructions show that gene innovation showed two peaks. Gene groups with the largest contribution to genomic change included extracellular proteins, transcription factors, as well as ones linked to the coordination of nutrient uptake with growth, highlighting the transition to a sessile osmotrophic feeding strategy and subsequent lifestyle evolution as important elements of early fungal evolution. Taken together, this work provided a highly resolved genome-wide catalogue of gene family changes across fungal evolution. This suggests that the genome of pre-fungal ancestors may have been transformed into the archetypal fungal genome by a combination of gradual gene loss, turnover and two large duplication events rather than by abrupt changes, and consequently, that the taxonomically defined fungal kingdom does not represent a genomically uniform assemblage of extant species characterized by diagnostic synapomorphies.

## Introduction

The evolutionary diversification of lineages into clades of various sizes, some highly successful and species-rich while others less so, is an outcome of complex interactions between ecological opportunity, changing biotic and abiotic environments and the genetic makeup of the organisms representing the lineage. Inferring the genomic footprint of the emergence of kingdom-level clades and, conversely, innovations that underlie their evolutionary success have been hard to assess. Recently, comparative genomic approaches that allow ancestral genome content to be inferred based on extant genomes revealed that complex mechanisms, including novel and co-opted genes as well as gene loss^1–3^ contributed to the origins various clades, including pro-^4^ and eukaryotes^5^, metazoans^1,6^ or land plants (Embryophytes^2^). These patterns are consistent with proposed general trends of genome evolution^7^, however, whether they are universal across the tree of life is unknown.

Fungi are one of the evolutionarily most successful groups. They not only exhibit an extreme diversity in morphological traits, but also in ecological functions, which make them key players in the ecosystem as symbionts, parasites or saprobes, among others. Kingdom Fungi, defined in the broad sense to include Rozellomycota and Microsporidia (also known as Opisthosporidia) and Aphelidomycota^8^, encompasses at least 10 phylum-level clades, with most of its phylogenetic diversity found in early-diverging fungi (EDF), an informal group comprising Rozello-, Aphelido-, Chytridio-, Sanchytrio-, Blastocladio-, Olpidio-, Zoopago- and Mucoromycota as well as Microsporidia^9,10^. Irrespective of significant flux in the taxonomic definition of Fungi^11^, how the earliest fungal ancestors are characterised and how they evolved from a unicellular opisthokont ancestor has been the subject of intense research. Reconstructions of the last universal fungal ancestor (LUFA) became more and more nuanced with the inclusion of newly discovered and/or classified early diverging lineages^8,10,12–16^. These results contributed to our current understanding of LUFA as a unicellular phagotrophic parasite of microalgae, possessing motile cell stages with amoeboid and flagellar motility and a chitinous cell wall, at least in part of its life cycle^9,15^. In contrast, derived fungi (i.e., Dikarya) are sessile terrestrial osmotrophs that grow septate hyphae, and their cells are covered by a rigid cell wall throughout their life cycle. However, how LUFA and subsequent ancestors evolved modern fungal traits is poorly known, however, and systematic analyses of the genomic changes during this transformation are missing.

In this study we reconstructed gene family evolution in the fungal kingdom using 123 whole genomes and analysed temporal and functional trends in the genomic changes we inferred. We find that early diverging fungi are genetically intermediate between pre-fungal protists and Dikarya and that a gradual shedding of ancient protist gene families happened in parallel with the emergence of fungal novelties and expansion of pre-existing families in Fungi. The tempo and mode of gene turnover and innovation outlines major genomic trends and picture an episodic emergence of modern fungal traits, including multiple waves of gene family expansion and contraction related to key fungal traits. Our results reveal that taxonomically defined fungi don’t match with ‘genomic fungi’ and that early fungal evolution has been highly episodic with continuous gene turnover.

## Results and Discussion

### Early diverging fungi are intermediate between protists and the Dikarya

To obtain a global perspective on gene content differences between fungi and related opisthokonts, we first clustered protein sequences of 123 species into homologous protein groups (HGs, see Methods). For this, we selected representatives of all, but one currently accepted fungal phyla (including Rozellomycota and Microsporidia, except Sanchythriomycota), as well as 17 Holozoa, Amoebozoa and Heterolobosea species as outgroup (Supplementary Data 1). We inferred a species phylogeny by maximum likelihood analysis of a supermatrix of 272 single-copy orthologs (Fig. 1a, Supplementary Fig. 1). Our phylogeny is highly supported and is largely congruent with phylogenies from recent studies^9,15,17–19^.

**Figure 1.**
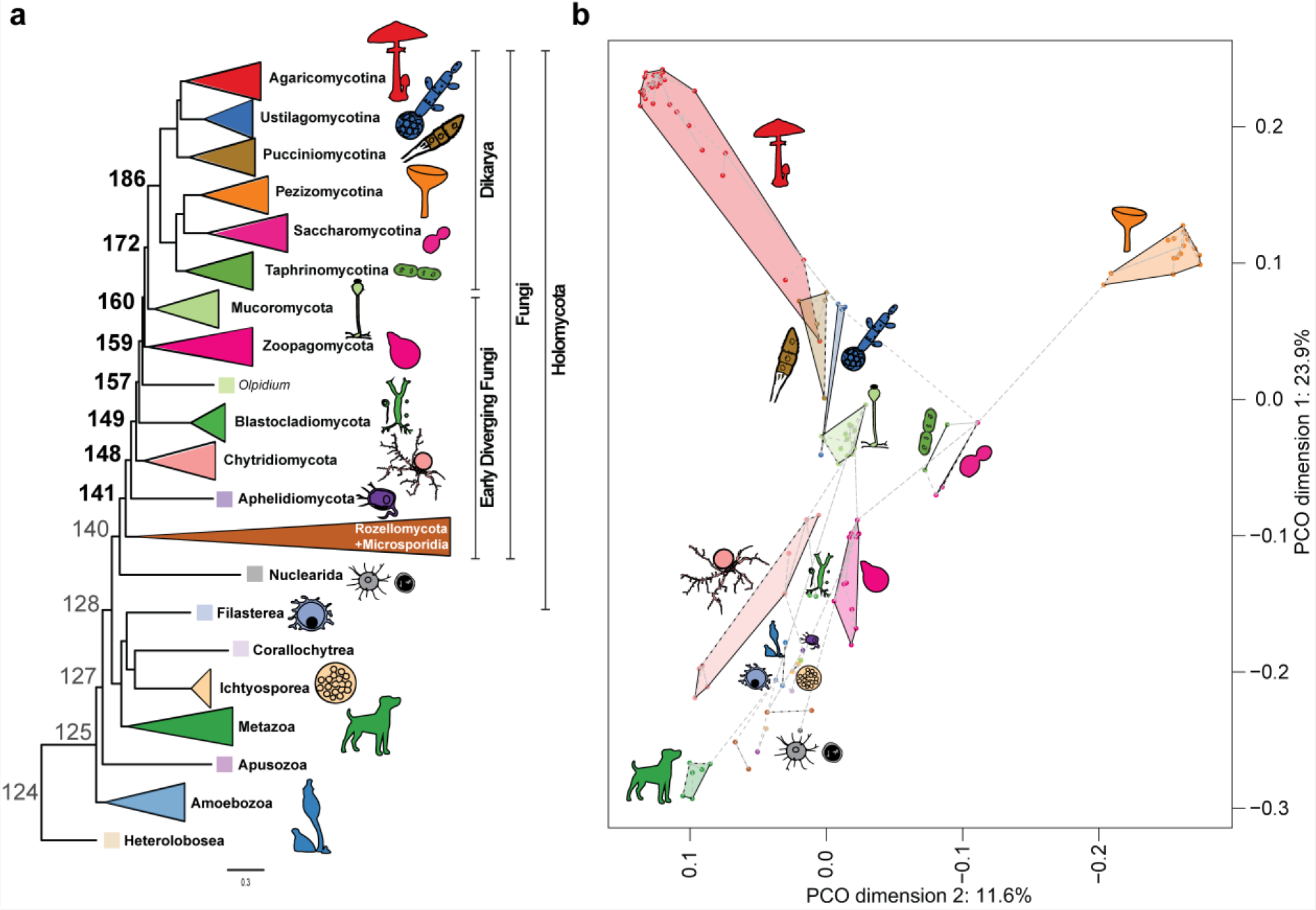
Intermediate position of early diverging fungi between protists and Dikarya. **a)** Schematic representation of our species tree showing the taxonomic clades and node numbers. Node names in bold are the fungal backbone nodes. See Supplementary Fig. 1 for the complete species tree and support values. **b)** Principal coordinate analysis (PCoA) and minimum spanning tree based on the presence/absence data of 9,993 HGs. Microsporidia species were removed from this analysis for better visualisation.

Based on the HG membership data, a Principal Coordinate Analysis resolved phyla into distinct groups, indicating that gene content divergence correlates well with phylogeny (Fig. 1b). Early diverging fungi, especially chytrids and aphelids, instead of occupying distinct positions in the space, grouped with non-fungal protists, with the Blastocladio- and Zoopagomycota being transitional towards the Mucoromycota+Dikarya (Fig. 1b). In contrast, the Dikarya branched into two groups in a tree-like pattern corresponding to the Asco- and Basidiomycota, with secondarily reduced yeast-like lineages of both phyla falling closer to each other. The intermediate placement of early diverging fungi in the genomic space is consistent with a number of phenotypic (amoeboids, flagellated zoospores), ecological (parasites, phagotrophic feeding), biochemical and genetic similarities (see^20,21^). Some of these have already been studied in detail, like the distribution of flagellar genes^22^, cytoskeletal complexity^23^, class V-VII Chitin synthases^22^, cell-cycle system^24^, WASH complex^13^ or cobalamin utilisation^25^. However, whether protist genes are ubiquitous in EDF has not been systematically investigated. Based on these observations we asked whether the presence of protist-like genes is a broad genomic trend in EDF, and if so, how were these replaced by fungal novelties during the evolution of the fungal kingdom.

### Gradual turnover of protist heritage and synapomorphies in fungi

By systematically searching for HGs with >70% conservation in at least one taxonomic group (Supplementary Data 1), present in the Holomycota but absent from Dikarya, we identified 540 families (Fig. 2, Supplementary Data 2). This indicates that EDF possesses a considerable number of HGs shared with protists, considerably more than previous anecdotal evidence suggested. For example, LUFA lost 41 of these, but possessed 499 HGs that are conserved in protists but are missing in Dikarya (Fig. 2). Analysis of the extinction dynamics of the 540 HGs showed that they were lost in a stepwise manner, with the largest loss events, 76, 64 and 239 HGs inferred in nodes where Blastocladiomycota, Zoopagomycota and Dikarya, respectively, split from the backbone.

**Figure 2.**
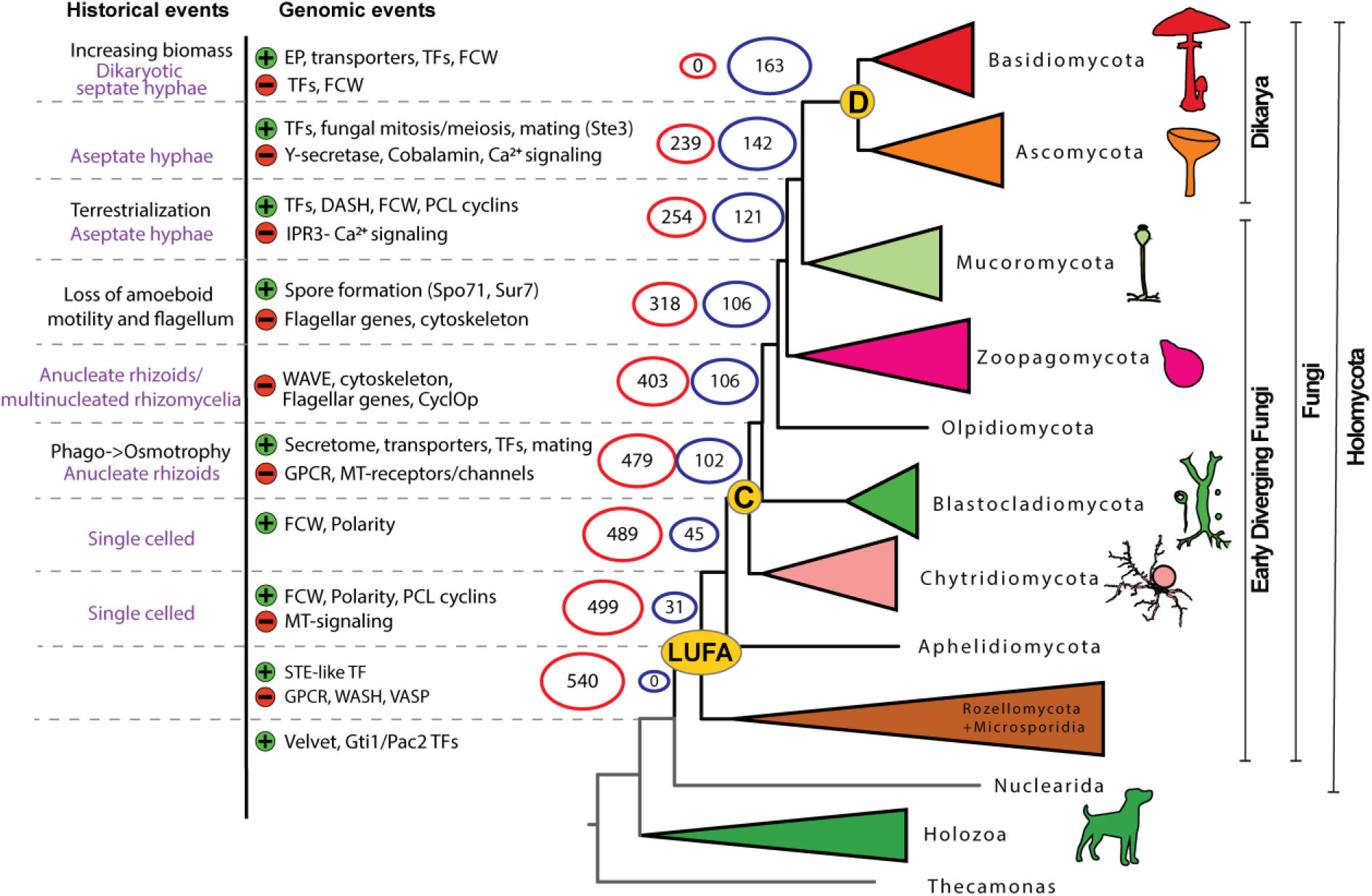
Changes in homologous groups presence/absence during fungal evolution. Historical events according to Berbee et al.^20^, Kiss et al.^34^, probable cellular complexity (with purple) and noteworthy genomic events (green “+” gain/expansion, red “-” loss/contraction) that we detected are at corresponding nodes. Numbers at nodes indicate the number of conserved protist-specific (red) and fungal novelty (blue) HGs present at each node. Abbreviations are as follows: LUFA: Last Universal Fungal Ancestor, C: MRCA of Chytridiomycota and derived fungi, D: MRCA of Dikarya, EP: extracellular proteins, MT-signalling: Metazoa-type signalling (e.g. Ca^2+^ signalling, EGFR), GPCR: G-protein coupled receptors, WASP: Wiskott-Aldrich syndrome protein, VASP: vasodilator-stimulated phosphoprotein, FCW: Fungal cell wall, PCWDE: plant cell wall degrading enzymes, TFs: transcription factors, E2F: E2F type TF, DASH: fungal specific part of the kinetochore complex.

The 540 HGs included all gene families previously reported to be lost in EDF: subunits of WASH and VASP (lost in LUFA), WAVE^14^, the CyclOP^26^ light sensing system (lost after Blastocladiomycota), flagellar genes^22^ (lost across the branching of Chytridiomycota, Blastocladiomycota, and *Olpidium*, see Supplementary Data 3), cobalamin synthesis proteins^25^, the replacement of the E2F cell cycle regulator by the SBF complex^24^. However, different taxon sampling updated the inferred time point of these loss events in several cases. Moreover, these events covered only a minority (73 HGs, 13.5%) of the 540 HGs. To broadly understand the functions of gene families lost in EDF, we performed Gene Ontology enrichment analyses (GO) in the 540 HGs, for each backbone node (Fig. 2, Supplementary Data 2). It revealed the significant overrepresentation of GO terms related to Ca^2+^-binding proteins and members of IP3, DAG, and Ca^2+^ signalling pathways.

We inferred both a copy number reduction and complete loss of HGs within these pathways (Supplementary Fig. 2-3). Shared ancestry of Ca^2+^ signalling in Fungi and Metazoa was known, despite of the remarkable differences among them^27^, but our results revealed that the full complement of these pathways have been retained in fungi until the Mucoromycota-Dikarya MRCA. Our data set also showed that the gamma-secretase complex, which is involved in regulated intramembrane proteolysis for various developmental and signalling processes and was thought to be absent in fungi^28^, in fact, was lost only in Dikarya, Olpidium, and the RM clade. Interestingly, several HGs associated with ubiquitination have also been lost during fungal evolution. Other functions highlighted by GO correspond to families annotated in extant holozoa as mechano- and voltage-sensitive channels, receptors (EGF, GPCRs) and mechanical or visual perception (Supplementary Data 2). Metazoan annotations of these families are the most precise that exist, but may be hardly informative as to the functions of these families in pre-fungal protists and EDF.

We also looked at core fungal novelties, defined here as HGs that evolved in one of the early fungal ancestors and are highly conserved (≥70%) in descendant lineages. Previous studies reported a shortage of fungal synapomorphies^11^, so we here tested if our genome-wide dataset can recover gene families specific to and conserved within fungi. We identified 163 HGs, considerably less than recent studies have, in animals and green plants (using 95% conservation threshold^1^). This suggests that novel core gene families are less prevalent in fungi, possible because fungi comprise several highly reduced clades (e.g. yeast lineages^30^, Microsporidia). Nevertheless, the 163 families originated across multiple nodes, with a larger grouping at the split of the Chytridiomycota (n=57, 35%, Node149, Fig. 2), suggesting that this node has seen key transitions in genome evolution (see 4^th^ chapter). Conserved domain analysis of these 163 HGs revealed that 72 contain protein domains that are predominantly (≥99%) found in fungi, corroborating them as real fungal novelties. These comprise functions in spore formation (e.g. Spo71 and Sur7 of *S. cerevisiae*), mating (e.g. Ste3, Prm1 and Rsc7/Swp82 of SWI/SNF complex of *S. cerevisiae*) or cell polarity (e.g. Spa2/Sph1, SOG2, of *S. cerevisiae*), among others (Supplementary Data 4a). We also detected families related to the cytoskeleton, the fungal cell wall (e.g. the Kre9/Knh1 family), intracellular trafficking, transporters and fungal-type mitosis/meiosis. We found a significant enrichment of transcription factors among novel HGs (Fisher’s exact test, P-value=0.002), including the origins of APSES, Copper fist, Opi1 and Fungal trans 2 families (see 4th chapter). Notably, the Velvet, Gti1/Pac2 and STE-like TF families, which are thought to be fungal specific, had representatives in early-diverging holozoans (*Capsaspora, Salpingoeca* and *Corallochytrium*) or in *Fonticula*. Using a domain-based search logic, we identified 186 HGs comprising fungal specific domains and contained functions relevant to sporulation, mating, intracellular transport, environmental sensing, chromatin remodelling, among others (Supplementary Data 4b). Finally, HGs containing unannotated proteins are prevalent (17.8% of 163 and 14.5% of 186 HGs) among core fungal novelties, highlighting the understudied status of fungal-specific genes.

Taken together, our analyses revealed that early-diverging fungi possess a large number of HGs shared specifically with protists, and that these were gradually lost during evolution. The broad conservation of protist HGs may explain the morphological and genetic similarities between EDF and protists reported in previous studies^16,23,24,31,32^ and also clarifies why the genome content of fungi reflects that of ancestral opisthokonts more than metazoan gene content does^33^. However, our data revealed that the retention of protist genes is not restricted to certain gene families, but is a genome-wide trend in early-diverging fungi. At the same time, we identified several conserved gene families that originated in early fungi and were mostly conserved afterwards, as well as 186 less conserved families containing fungal-specific domains. Although most of these can’t be considered synapomorphies in the strict sense^11^, due to their imperfect conservation, this indicates that beyond losses, considerable novelty has also emerged in early fungal ancestors.

### An interplay of gene loss and major bursts of gene duplication characterise fungal evolution

To obtain a detailed picture on gene repertoire changes and test if evolution is gradual or rather episodic, as some theories predict^35^, we reconstructed gene gain and loss dynamics for all HGs containing at least four proteins. Our reconstructions provide information on which genes were duplicated and lost in each of the HGs and at which branches of the phylogenetic tree (Fig. 3). For example, LUFA was inferred to have had 12,761 genes, gained 913 and lost 295 compared to its immediate ancestor, corresponding to a moderate net expansion. Reconstructed ancestral proteome sizes ranged from 12,761 protein-coding genes in LUFA to 14,891 in the MRCA of Zoopagomycota and derived fungi. If we consider net changes (duplications minus losses), it seems that most of the genomes of early fungal ancestors contracted (Fig. 3), with two exceptions. The first is the split of chytrids, while the second expanding ancestor is the MRCA of Zoopagomycota and derived fungi, which is inferred to have expanded by 791 genes (1,114 gains, 323 losses).

**Figure 3.**
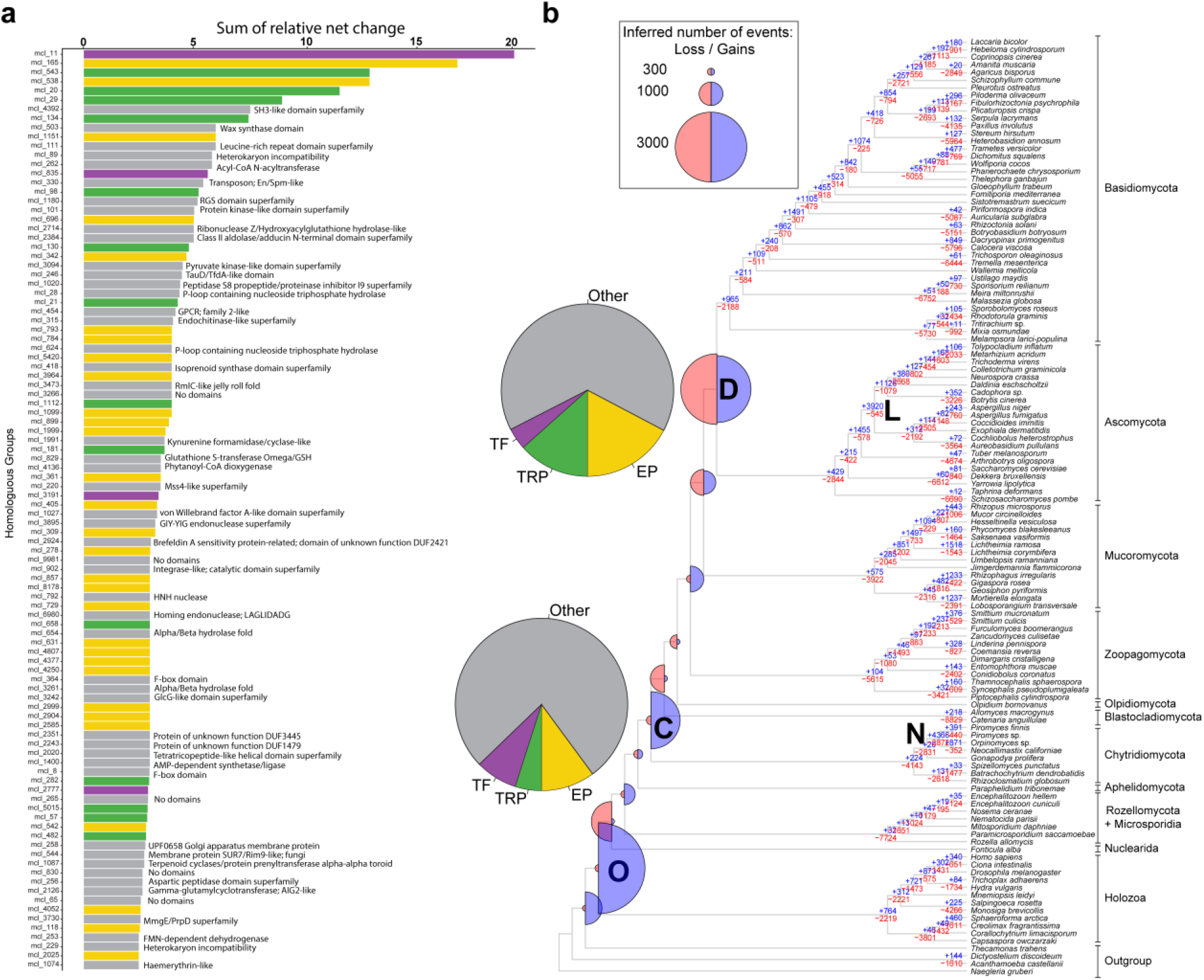
A global view of gene gains, duplication and loss across fungal evolution. **a)** The top 100 most dynamically changing HGs, coloured by functional categories (TF: transcription factors, TRP: transporters, EP: extracellular proteins). InterPro annotations are shown only for the ‘Other’ category. **b)** Gene duplication and loss history in the fungal kingdom. Blue (expansion) and red (contraction) half-circles indicate the inferred gains and losses at each node, while blue (gain) and red (loss) numbers along the backbone depict the number of gain and loss events. Note that the ratio of blue and red circles is proportional to the amount of net innovation and turnover in a given node. Letters in bold represent the five largest bursts of duplication that comprised more than 2,000 gain events. Pie charts, next to the nodes D and C represent the contribution of four functional categories to gene gains. Clade names are abbreviated as O: Opisthokonta MRCA, C: MRCA of Chytridiomycota and derived fungi, D: Dikarya MRCA, N: Neocallimastigomycotina, L: Leotiomyceta. Data for the first two nodes are not shown as they mostly contain pan-eukaryotic genes.

We find that gene duplication has been highly episodic, with five large burst events, whereas losses were ubiquitous in fungal backbone nodes. Of the five bursts of gene duplication two were inferred along the backbone of the fungal tree (nodes C and D on Fig. 3), two in phylum-level clades (N-Neocallimastigomycota and L-Leotiomyceta) and one in the Opisthokonta MRCA (O). From the two large duplication events we inferred along the fungal backbone, the first, in the MRCA of chytrids and other fungi (C), was coupled with limited gene loss (2,455 gains, 382 losses), whereas the second, in the Dikarya (D), coupled with more loss (2,875 gene gains, 3,036 losses), suggesting again, it was rather a turnover event. In general, the relative constancy of ancestral proteome sizes, in joint consideration with the losses and pulses of gene duplication indicates high gene turnover in early fungal evolution. Thus, while proteome sizes changed moderately, gene content has undergone significant changes.

Duplication events C and D contained a qualitatively similar functional signal, with an enrichment of terms related to extracellular functions (e.g. GO:0005576), transmembrane transport (e.g. GO:0055085), cellulose binding (GO:0030248) and the fungal cell wall (e.g. GO:0016977). These may correspond to the transition from phagotrophy to osmotrophy and the chitinous cell wall, respectively, and reflect the improvement of extracellular digestive functions, as adaptations to the existence of increasingly complex and abundant plant material^20,36^. A significant enrichment of transcription factors (TF) was detected among duplicated genes across multiple nodes (Supplementary Data 5), consistent with results of novel core families (see above). Based on GO results, we identified an expansion of cyclins that transcends multiple nodes in early fungi (Supplementary Fig. 4, Supplementary Data 5). In extant fungi, PCL cyclins coordinate the cell cycle with polarised growth in response to nutritional cues (phosphate, amino acid and glycogen metabolism)^37,38^, thus their diversification might have provided the basis of evolving sophisticated coordination of nutrient supply and growth. We detected further two bursts of duplication within fungi, one in the Leotiomyceta and one in the Neocallimastigomycota. These may correspond to periods of intense lineage-specific innovation which, in the latter clade, probably also reflects massive gene gains through HGT from bacteria^39^.

### Episodic genetic changes and major fungal innovations

We were next interested if the functional profile of inferred bursts of duplication correspond to hypothesised phenotypic changes in early fungal ancestors^20^. The most dynamically changing (expanding and contracting) gene families and corresponding functions were identified by ranking families by their summed expansion/contraction dynamics across eight nodes from LUFA to the MRCA of Dikarya (Supplementary Data 6). Among the 100 most dynamically expanding families, 51 contained extracellular proteins (mostly CAZymes), transporters or transcription factors (Fig. 3a). The remaining families included diverse functions, such as GPCRs, heterokaryon incompatibility genes, protein kinases and endochitinases, among others (Supplementary Data 6). When we parsed these figures in the context of the two large duplication events, we found that 22.9% of the chytrid and 34.7% of the Dikarya duplication bursts were related to extracellular functions, transporters and transcription factors, respectively (Fig. 3b). The importance of these gene families for genomic changes was also confirmed by functional enrichment analyses of HG duplication data (Supplementary Data 5). Based on these observations, we below scrutinise extracellular, transporter and transcription factor families in more detail.

Mapping of families with predicted extracellular localisation (hereafter extracellular proteins, EP, for short) revealed a two-stage duplication dynamics, with large expansions in the MRCA of chytrids and rest of the fungi (from 441 to 653 genes) and in that of Dikarya (from 806 to 1,167 genes; Supplementary Fig. 5). In the first event, EP expansion was driven mainly by CAZYmes, whereas other extracellular functions, including proteases, made a more modest contribution. These expansions resulted in a shift in the CAZYme and SSP content of the EP from 34% in LUFA to 51% in the Dikarya MRCA. Within EPs PCWDE families follow similar patterns, but with a larger expansion at the MRCA of Dikarya (Supplementary Fig. 6). In contrast to PCWDEs, FCW families showed proportionally more diversification in the chytrids than in the Dikarya, and large turnover in the Dikarya MRCA (Supplementary Fig. 7). The diversification of EP, especially PCWDEs in the ancestor of chytrids and other fungi, suggests their role in the transition from phagotrophy to osmotrophy, while expansion of FCW-related families here may have contributed to the evolution of the fungal cell wall. The second burst, in the Dikarya, may be concomitant with the radiation of plant lignocellulose degrading lineages, possibly in response to the onset of the radiation of land plants^36^. In the ancestors of Blastocladiomycota, Olpidiomycota and Zoopagomycota and other fungi the lack of PWCDE expansion and a moderate expansion of proteases may be explained with the primarily non-plant-based nutrition of these clades.

For cell surface transmembrane transporters, we identified 253 HGs based on characteristic domains and subcellular localization (Supplementary Data 7). Similar to CAZymes, transporters show a two-stage expansion, with the highest duplication rate inferred in the Dikarya MRCA (Supplementary Fig. 8). Of the three largest transporter families, we found that ABC transporters and P-type ATPases showed contraction, whereas the Major Facilitator Superfamily underwent an extreme expansion in fungi. This makes sense considering that only the latter is able to transport a high variety of small molecules, including sugars, peptides and lipids^40^, whereas ABC transporters and P-type ATPases are primarily exporters^41^ and specific for cations^42^, respectively. Given the role of transporters in osmotrophic nutrition^43^, the correlated diversification of transporters with CAZymes and the inferred continuous increase of copy numbers in ancient fungal genomes suggest their importance in the refinement of fungal-type heterotrophy.

We identified 657 transcription factor (TF) HGs based on domain content and classified them into 52 putative TF families (Supplementary Data 8) following the work of Sheleste^44^ and Mendoza et al.^45^. Mapping of these 657 HGs revealed a constant change in TFome during early fungal evolution. Looking at broad copy number dynamics, early nodes, such as the splits of chytrids (193 gains 3 losses) and Zoopagomycota (168 gains 5 losses) are almost exclusively dominated by gains with barely any losses (Fig. 4) This implies that the TFomes of these clades are similar to those of protists, as suggested by de Mendoza et al.^45^. In contrast, later evolution of EDF is characterised by the high turnover of TFs: at MRCA of Mucoromycota and derived fungi we inferred 87 gains and 43 losses while in the Dikarya MRCA we detected 117 gains 184 losses. Losses in the Dikarya MRCA affected several families (e.g. bZIP, Myb, CBF-NFYA, GATA, Homeobox), and included the complete disappearance of E2F TDP, T-Box, and Tub families. Along the backbone nodes TFome diversity (based on Shannon index) increased, with the highest number of ancestral TFs inferred in the MRCA of Mucoromycota and Dikarya (784 TFs; Fig. 4a). In line with this, extant Mucoromycota contain the most TFs among fungi, with Shannon-based diversities similar to those of EDFs (Fig. 4b). The TFomes of extant Dikarya, especially Ascomycota species show similar sizes but lower diversity, due partly to the expansion of the Fungal trans 2 and Zn cluster TF families, and partly to losses of certain families in the Dikarya.

**Figure 4.**
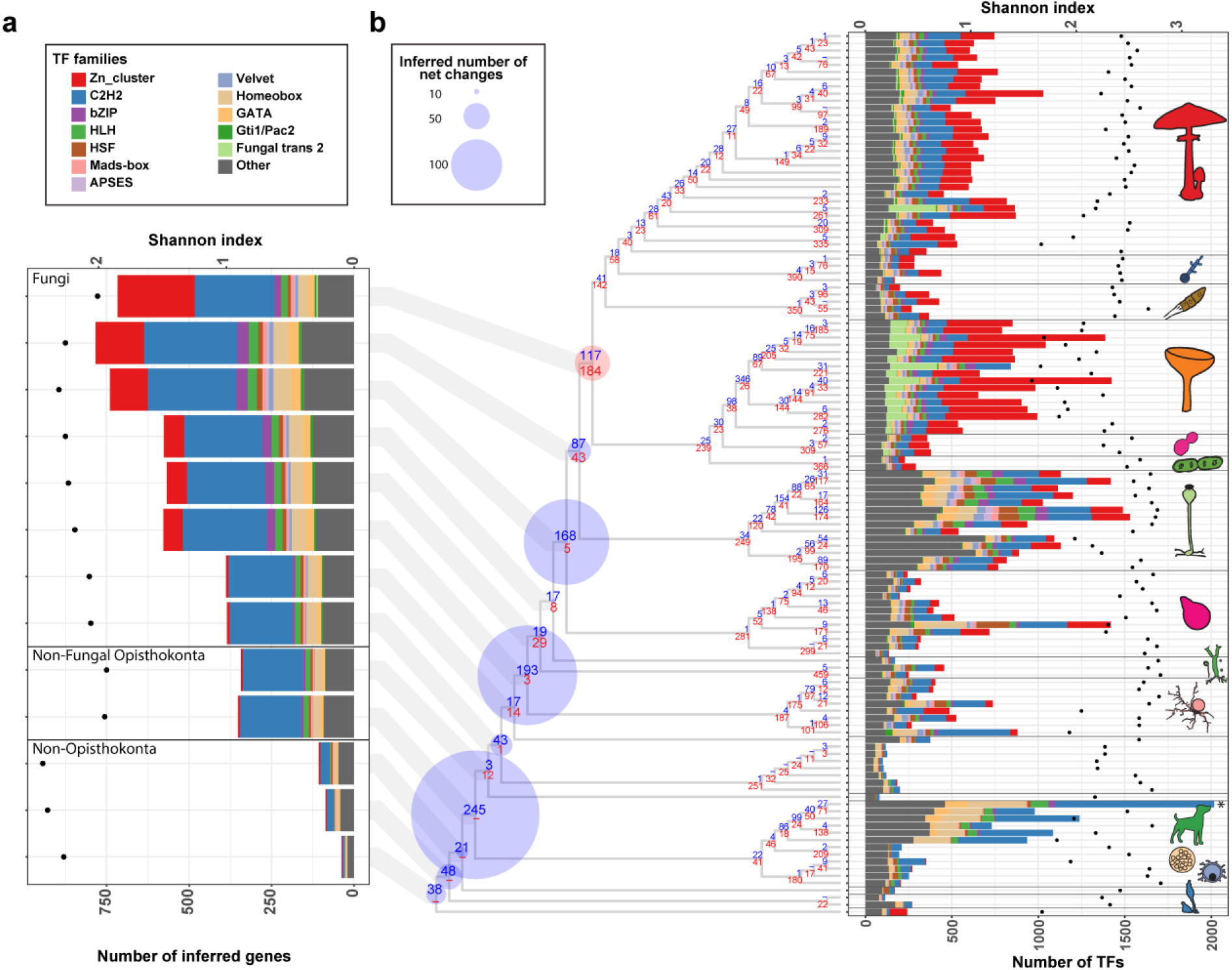
Dynamic turnover of transcription factor families. **a)** Inferred ancestral size of transcription factor families in fungal backbone nodes. Only the 12 families that showed the highest dynamics in early fungal evolution were coloured differently (see main text and Supplementary Data 8). **b)** Inferred net changes based on the 657 TF HGs mapped to the species tree (left). Blue (expansion) and red (contraction) circles indicate the inferred net changes (gains minus losses) at each node, while blue (gain) and red (loss) numbers along the backbone depict the number of gain and loss events. The stacked bar chart (right) represents the size of TF families, using the same colouring scheme as in (**a**). For visualisation, the *Homo sapiens* column (*) was scaled down proportionally from 5,262. The Shannon index of TF-ome diversity for inferred ancestral (a) and extant (b) species is shown with dots.

Inferred copy number of TFs in fungal ancestors, outlined three ‘epochs’, within each of which ancestral TF repertoires seemed to be relatively constant (Fig. 4a). Transitions between these epochs are concominant with remarkable historical events, like emergence of—mostly anucleate—rhizoids and osmotrophy, or that of terrestrialization and of aseptate hyphae (Fig. 2 and 4a).

Ranking of TF families based on the cumulative net change along the fungal backbone (Supplementary Data 8) revealed that The most dynamically changing TF families included both pan-eukaryotic (e.g. C2H2-like, bZIP, HLH, HSF, Homeobox, GATA) and predominantly fungal TFs (e.g. Zn cluster, Fungal trans 2, APSES, Velvet and Gti1/Pac2), irrespective of their raw copy numbers. Several families expanded considerably more than the whole proteome, indicating that their copy numbers are decoupled from proteome size. For example, while the proteome size of the MRCA of Chytrids and derived fungi increased by 16.2% relative to the preceding node, the Zn cluster and bZIP families expanded by 738% and 325%, respectively; suggesting expansion driven by adaptation. Interestingly, C2H2 TFs underwent a massive expansion in the Opisthokonta ancestor. This family is highly diverse in both Holozoa and the Holomycota, with clade specific contractions in secondarily simplified clades, such as yeast-like Dikarya (Supplementary Fig. 9).

Taken together, the evolutionary dynamics of transcription factors differs from genome-wide patterns, in that, albeit also episodic, showed expansions in different nodes. Nevertheless, these data suggest that broad rewiring of fungal gene regulatory networks has transcended multiple ancestors in EDF.

## Conclusions

The origins of highly diverse clades across the tree of life are remarkable evolutionary events, but are they remarkable from a genomic perspective or because we attach taxonomic definitions, such as a kingdom, to some of them? Based on systematic analyses we inferred that protein coding gene content has changed drastically during early fungal evolution. However, in contrast to animals and plants^2^, this has not happened abruptly at the taxonomic limit of the kingdom, even if competing taxonomic circumscriptions of Fungi are considered. Rather, we found a significant retention and stepwise loss of protist genes in early diverging fungi, combined with the episodic expansion of novel and ancient gene families. We also identified major functional trends and the most dynamically changing gene families, which allowed us to relate genomic changes to trait evolution during the early evolution of fungi.

Our systematic analyses revealed that early-diverging fungi retained hundreds of protist gene families that are missing in the Dikarya. These were lost gradually or replaced by fungal-specific genes (e.g. E2F cell cycle regulator by SBF^24^) during fungal evolution. We think this explains why EDF gravitate towards protists in gene content-based analyses (See Fig. 1) and provides a genome-wide explanation for sporadic evidence for the similarity of EDFs to protists from analyses of single genes^24,25^, individual genomes^33^ or TF-omes^45^. At the same time, we detected a limited number of gene families that could be considered synapomorphic for fungi, in agreement with previous conjectures on the lack of synapomorphies in the kingdom^11,22^.

The retention of protist genes and the lack of clear synapomorphies for fungi blur lines between fungi and related protists and may explain why the taxonomic limits of fungi have been challenging to define and are still a matter of debate^14^. In other words, taxonomically defined Fungi don’t match with any clade we could define, based on gene content, as ‘genomic Fungi’, a situation that would not change with the exclusion of Opisthosporidia from fungi. Defining ‘genomic Fungi’ is also complicated by patterns of gene turnover, nevertheless, gene content (Fig. 1), turnover rates (Fig. 2) and novel gene families suggest that, if any fungal clade, the Dikarya comprises species with a broader set of signature gene families.

Gene loss has been recently named as a dominant mechanism of metazoan^3,46,47^ and plant evolution^2^. Similar patterns were often observed in ancestral genome reconstructions, which gave rise to the hypothesis of genome evolution comprising periods of genome expansion resulting in complex ancestral genomes, followed by genome streamlining via gene loss^35^. In contrast, early fungal evolution may be better viewed as a series of turnover events, in which gene loss aided the gradual shedding of protist traits whereas innovation was probably fueled by two bursts and continuous small-scale duplication of genes.

Functional analysis of gene duplications and losses outlined major functional trends in the evolution of early fungal genomes. Remarkable lost or contracting gene groups were related to phagocytosis, the flagellum, cell cycle regulation and signalling (Fig. 3), among others. For highly duplicated gene groups, a dominant functional signal was related to extracellular proteins (e.g. secreted CAZymes) and transporters which can be linked to the transition from phagotrophy to osmotrophy and the subsequent sophistication of extracellular digestive and uptake mechanisms^11,14,15,20,34,48^. These also explained a considerable portion of the duplication events in the chytrid and Dikarya ancestors, though it should be noted that a myriad other functions were also represented among the most expanding families. For example, transcription factors emerged as one of the most dynamically changing gene groups, suggesting a broad rewiring of gene regulatory mechanisms in early fungi. Another group of dynamically changing regulators were PCL-type cyclins, which link nutritional status of fungal cells with the cell cycle and thus the expansion of this family is perhaps related to balancing nutrient assimilation and growth, an important regulatory mechanism for hyphal osmotrophs. Our genome-wide catalogue of changes revealed a large array of other functions as well, many of which cannot currently be easily linked to phenotypes or ecological functions due to the paucity of knowledge on gene function.

Taken together, this study reconstructed the history of genomic change in the fungal kingdom at unprecedented detail and provides a resource for further analyses of fungal genome evolution, also at smaller evolutionary time scales. We conclude that, albeit sharp genetic changes at the border of the fungal kingdom were not inferred, a large turnover of protistan genes and a gradual emergence of fungal novelties in early fungal evolution portray a clear genetic roadmap for the emergence of modern fungi from a unicellular algal parasite ancestor.

## Methods

### Dataset assembly and clustering of proteins

To evenly sample clades in the fungal kingdom, we sampled 106 fungal species, consisting of proteomes from all known phylum level clades, with the exception of Sanchytriomycota, which were not published prior to our data collection. In addition, as outgroups, 17 non-fungal species were sampled to represent Heterolobosea, Amoebozoa, Holozoa, and Nucleariids. Proteome sequences were downloaded from the JGI Genome Portal and NCBI/Ensembl (before July 2021;^49,50^). All-vs-all similarity searches of the 123 proteomes were performed with MMseqs2^51^ using three iterations, with sensitivity set to 6.5, max-seqs to 15,000, e-profile set to 0.001, a preliminary coverage threshold set to 0.2, and an e-value threshold set to 1e^-4^. We then performed an asymmetric coverage filtering (requiring 20% coverage for the longer and 80% for the shorter protein) and Markov clustering with an inflation parameter 2.0^52^ as described previously^53^. After clustering, we removed contaminating proteins from gene families following the logic of Richter et al.^47^. Further, to achieve better completeness of clusters without increasing noise, we merged clusters based on similarity, using the all vs all output of MMSeqs and the results of HMM search between the consensus sequence and HMM profiles of clusters. Based on the MMSeqs output file, the connection intensity of clusters (i.e a network) was constructed. These cluster networks were iteratively reduced by excluding the weakest nodes-sorted by the number of connections between the two clusters, normalised to cluster size-until the maximum diameter of a network was three. Of these cluster pairs, only those that achieved an e-value cut-off at least 1e^-10^ in the HMM search (http://hmmer.org/) with an asymmetric coverage of 75/20% (HMM profile and consensus, respectively) and whose match score reached at least 75% of the self-match were allowed to be merged. We called these merged filtered clusters of protein as homologous protein groups, briefly HGs.

### Species tree reconstruction

For species tree inference, marker genes were selected from four sources: 1) single-copy gene families based on the clustering mentioned above, 2) clusters that were single-copy after eliminating terminal duplications, 3) HMM-based search of BUSCO, and 4) HMM of JGI 1086 marker gene sets^54,55^ (https://github.com/1KFG/Phylogenomics_HMMs/tree/master/HMM/JGI_1086). For the latter two, only best hits were used for each species. Subsequently clusters were removed, where the average distance of amino acid (AA) alignment were too high (>=1.5; dist.ml with the model WAG^56^) and which showed paraphyletic clusters in the hierarchical clustering of AA distances. Multiple sequence alignments were inferred using PRANK v.170427^57^ and trimmed using TrimAL v.1.2^58^ (-strict). Trimmed MSAs shorter than 60 amino acid residues (AA) and containing < 30 species were discarded, leaving 272 single-copy clusters resulting 68,662 sites that were finally used for tree reconstruction. Phylogenetic inference was performed under maximum likelihood (ML) in IQ-TREE v1.6.12^59^ with ultrafast bootstrap^60^ (1000 replicates) based on the partitioned dataset of 272 clusters using the substitution model LG+G. More complex models (LG+C60+G) had no effect on the topology of early diverging lineages. We applied a constrained tree topology ((Allma1,Olpbor1),Ganpr1,Partr) in order to separate Aphelidomycota from Blastocladiomycota, which caused no significant change in log-likelihood values as assessed by the Shimodaira-Hasegawa (SH) test^61^ (deltaL=21.8, p-value = 0.356).

### Inference of genome-wide duplication and loss history of clusters

For gene tree reconstructions, gene families containing at least four proteins were aligned using the L-INS-I or auto (if the former was not applicable) algorithm MAFFT v7.313^62^ and trimmed with TrimAL (-gt 0.2). Gene trees were inferred in RAxmlHPC-PTHREADS-AVX2 8.2.12 under the PROTGAMMAWAG model and we estimated branch robustness using the SH-like support^63^. Rooting and gene-tree/species-tree reconciliation was performed with NOTUNG v2.9^64^ using an edge-weight threshold of 80.

Gene duplication and loss histories were inferred by mapping orthologous groups delimited on the basis of gene trees to the species tree using Dollo parsimony implemented in a modified version of COMPARE^30^. Detected gene gains could be de novo origination, duplication, horizontal gene transfer, or the result of undetectable distant homology, however in this study, we did not attempt to separate these events. A custom R script (https://github.com/zsmerenyi/compaRe) was used to visualise the mapping results utilising functions of the phytools, ape, tidyr and phangorn packages^56,65–68^.

### Annotation of homologous groups

For the evaluation of domain content and gene ontology (GO) terms of HGs, an InterProScan-5.47-82.0^69^ was performed on all 123 proteomes. A GO enrichment was carried out using Fisher’s exact test with the weight01Fisher algorithm of the topGO Bioconductor module^70^, and a p-value <0.05 was considered significant. For the enrichment analysis of 540 gene families lost across fungal evolution, the most complete, *Homo sapiens* GO list was used as a reference. For the GO enrichment analysis of HGs that underwent duplication or loss events along the fungal backbone, all species were used and frequencies of ortholog groups were taken into account for each node.

A uniform rule was used for further annotation of HGs: a HG was assigned to a group if >50% of its proteins were annotated with the same type of annotation (e.g., a specific domain, SSP, or extracellular localisation). For transcription factor (TF) family identification, sequence-specific DNA binding domains (DBDs) were selected based on literature mining. From the putative TF HGs (containing > 50% of a given DBD), we filtered out those containing domains characteristic of non-TF families or processes such as ribonucleases, metallopeptidases, chromatin remodelling, or splicing. Finally, we classified the HGs into TF families based on their DBD content, based on previous works^44,45^.

Extracellular protein identification was based on subcellular localization prediction by WoLFPSORT 0.2^71^ using the “fungi” option. Proteases, small secreted proteins (SSPs), and carbohydrate-active enzymes (CAZYmes) were predicted to further differentiate the extracellular protein-containing HGs. For proteases, the non-redundant database was downloaded from MEROPS (on 27, sept. 2022, from https://www.ebi.ac.uk/merops/download_list.shtmlmerops_scan.lib) and used as a query against the protein sequences of 123 species in a BLAST search, with 20% bidirectional coverage, 1e-5 e-value cutoff, keeping the best hit for each subject protein. Prediction of SSPs was performed using a modified version of the bioinformatics pipeline of Pellegrin et al.^72^, as follows. Proteins shorter than 300 amino acids were subjected to signal peptide prediction in SignalP (version 4.1^73^). Proteins containing a transmembrane helix-predicted by TMHMM version 2.0^74^-were excluded. For the prediction of CAZYmes, a HMM search was performed with dbCAN2, using dbCAN-HMM profiles (http://csbl.bmb.uga.edu/db CAN^75^) as queries. Subsequently, CAZYmes were classified according to the work of Hage & Rosso^76^. Fungal cell wall (FCW) and plant cell wall degrading enzymes (PCWDE) were based on the classification of CAZY families in previous works^77,78^.

Detection of transporters was based on the presence of characteristic InterPro (IPR) domain according to the work of Sahu et al.^79^ and plasma membrane localization, predicted by WoLFPSORT. Only those HGs were considered transporters that contained >50% of transporter-specific domains, and if plasma membrane localization (score >15) was the most likely within the HG.

### Identification of conserved and dynamically changing homologous groups

To obtain an overall picture of similarities and differences in the gene repertoire of the 123 species, a principal coordinate analysis (PCoA) was performed. For PCoA, a total of 9,993 homologous groups (HG) comprising at least four proteins and reaching ≥50% species from any clade, were used (see clades in Supplementary Data 1). A binary distance coefficient from the gene family presence/absence data was used, and a minimum spanning tree was superimposed on the distribution of species on the two principal coordinates (using packages of stat, ape and tidyverse^66–68^).

To identify conserved HGs, we defined eight groups, from which the ‘non-opisthokonta outgroups’ and ‘basal Holozoa’ are paraphyletic, while Metazoa, Chytridiomycota, Zoopagomycota, Mucoromycota, Pezizomycotina, and Agaricomycotina are all monophyletic (see Supplementary Data 1). Conservation of a HG was calculated as the proportion of the number of species with protein present to the total number of species in each of these groups. We considered HGs to be shared between EDF and protists if they had at least 70% conservation in any of the above groups, the HG emerged in or before the first Holomycota node and was missing in Dikarya.

For identifying fungal core novelties, we searched for HGs that emerged after node 141 (LUFA) and had >=70% conservation for all descendants of the node in which it emerged. This is similar to ‘novel core’ families investigated previously in animals^80^ and plants^2^, however, we chose a conservation threshold of 70%, due to the large number of secondarily simplified lineages (e.g. yeasts, Microsporidia) among fungi^81^.

To validate fungal core novelties and losses, the distribution of InterPro domains among high-ranking taxonomic groups (Viruses, Bacteria, Archaea, Chromalveolata, Chromista, Excavata, Plantae, Protists, Holomycota, Holozoa) was examined. InterPro annotated proteins from UniProtKB together with InterPro annotation (Release 2022_01), and the NCBI taxonomy database^82^ (in 05/2022) were downloaded. The distribution of the 37,834 IntePro domains among 10 taxonomic groups was calculated by normalising the total counts for each domain; based on altogether 230,895,644 proteins. We used this dataset to assess fungal dominance of a domain, for example a domain considered as fungal specific if 99% of uniprot hits come from Holomycota (see Supplementary Data 4). Also this proportion was used for mining HGs containing at least 75% of fungal-specific domain (Supplementary Data 4b).

To assess the dynamics of HG changes along the fungal backbone, the relative net change to the copy number of the preceding node was calculated and summed over across eight nodes from LUFA to the MRCA of Dikarya (see Supplementary Data 6 and 8).

## Acknowledgements

We appreciate the critical comments of Balázs Papp, and Eduard Ocaña-Pallarès on the earlier version of the manuscript. Attila Csikasz-Nagy is thanked for useful discussions on PCL cyclins. We are thankful to Mary Catherine Aime for permission to utilize unpublished genomic data of *Tritirachium sp*. This work was funded by the Momentum Program of the Hungarian Academy of Sciences (LP2019-13/2019) and by the European Research Council (Grant No. 758161) (both to LGN).

## Author contributions

L.G.N., K.K, J.S. J.W.S., Z.M., designed the research, K.K., N.S., X.B.L, B.B., and Z.M., performed Data collection and curation Z.M., performed the comparative analyses and bioinformatics. L.G.N., J.W.S., and Z.M. interpreted the results and wrote the paper. All authors have read and agreed to the published version of the manuscript.

## Legends for supplementary data

**Supplemental Data 1**. Taxon list of 123 species used for our analyses. BUSCO percentages were obtained with the database ‘eukaryote’.

**Supplementary Data 2a**. Conserved HGs shared between protists and early diverging fungi (EDF). 540 HGs which showed at least 70% conservation in any of the groups defined in Supplementary Data 1, emerged before node 140 (split of Nuclearia) and were lost along the backbone between nodes 141 (split of RM clade) and 186 (MRCA of Dikarya).

**Supplemental Data 2b**. Enriched functional categories among HGs (540) lost during fungal evolution. The matrix contains only the significantly enriched (p-value <= 0.05) terms for each backbone node of the tree. Abbreviation n.s. means not significant.

**Supplementary Data 3**. Flagellum-associated proteins. We used the list of flagellar proteins of *Chlamydomonas reinhardtii* from (http://jcb.rupress.org/content/170/1/103.full) as queries in MMSeqs reciprocal best hit searches in the proteomes of the examined species. Numbers in a matrix are -log10 transformed e-values. Species highlighted with green possess flagella.

**Supplementary Data 4a**. Novel core HGs in fungi. 163 HGs which showed at least 70% conservation and were gained along the fungal backbone (>= node 141 =< node 186).

**Supplementary Data 4b**. Homologous groups containing fungal-specific domain, and emerged in the backbone (>= node 141 =< node 186).

**Supplementary Data 5a**. TopGO functional enrichment for the emergence of orthogroups (duplication or de novo birth). Values in the matrix represent the p-value of Fisher’s exact test with weight01Fisher algorithm of topGO. Only values < 0.05 are shown. ‘n.s.’ means not significant.

**Supplementary Data 5b**. TopGO functional enrichment for the 540 orthogroups shared between protists and EDF. The node and clade names represent the last ancestor in which these orthogroups are predicted to be still present. Values in the matrix represent the p-value of Fisher’s exact test with weight01Fisher algorithm of topGO. Only values < 0.05 were visualised. ‘n.s.’ means not significant.

**Supplementary Data 6a**. The dynamics of expansion/contraction of HGs during early fungal evolution.

To assess the dynamics of HG changes along the fungal backbone, net change relative to the copy number of the preceding node was calculated and summed across eight nodes from LUFA to the MRCA of Dikarya. We ordered our table based on this score (‘Sum of relative net change’).

**Supplementary Data 6b**. Inferred gain, loss events and copy numbers of all HGs in the MRCA of Chytridiomycota and other fungi. 27.2% of gains are due to the gene families analysed.

**Supplementary Data 6c**. Inferred gain, loss events and copy numbers of all HGs in the MRCA of Dikarya. 40.7% of gains are attributable to the analysed gene families.

**Supplementary Data 7**. 253 identified transporter HGs, their classification and distribution among clades.

**Supplemental Data 8a**. Evolutionary dynamics of transcription factor families. Column “cumulative net change” means the sum of normalised net changes (gains minus losses) by the copy number of the parental node. The first 12 families (in bold) were visualised in Fig. 4.

**Supplementary Data 8b**. 657 identified TF HGs, their classification and distribution among clades.

