## Supplementary material for "Taxonomic vs genomic fungi: contrasting evolutionary loss of protistan genomic heritage and emergence of fungal novelties": Supplementary_Figs_2022.pdf

This PDF file includes:

Supplementary Figures 1 to 9
Captions for Supplementary Data 1 to 8

### Supplementary Figures

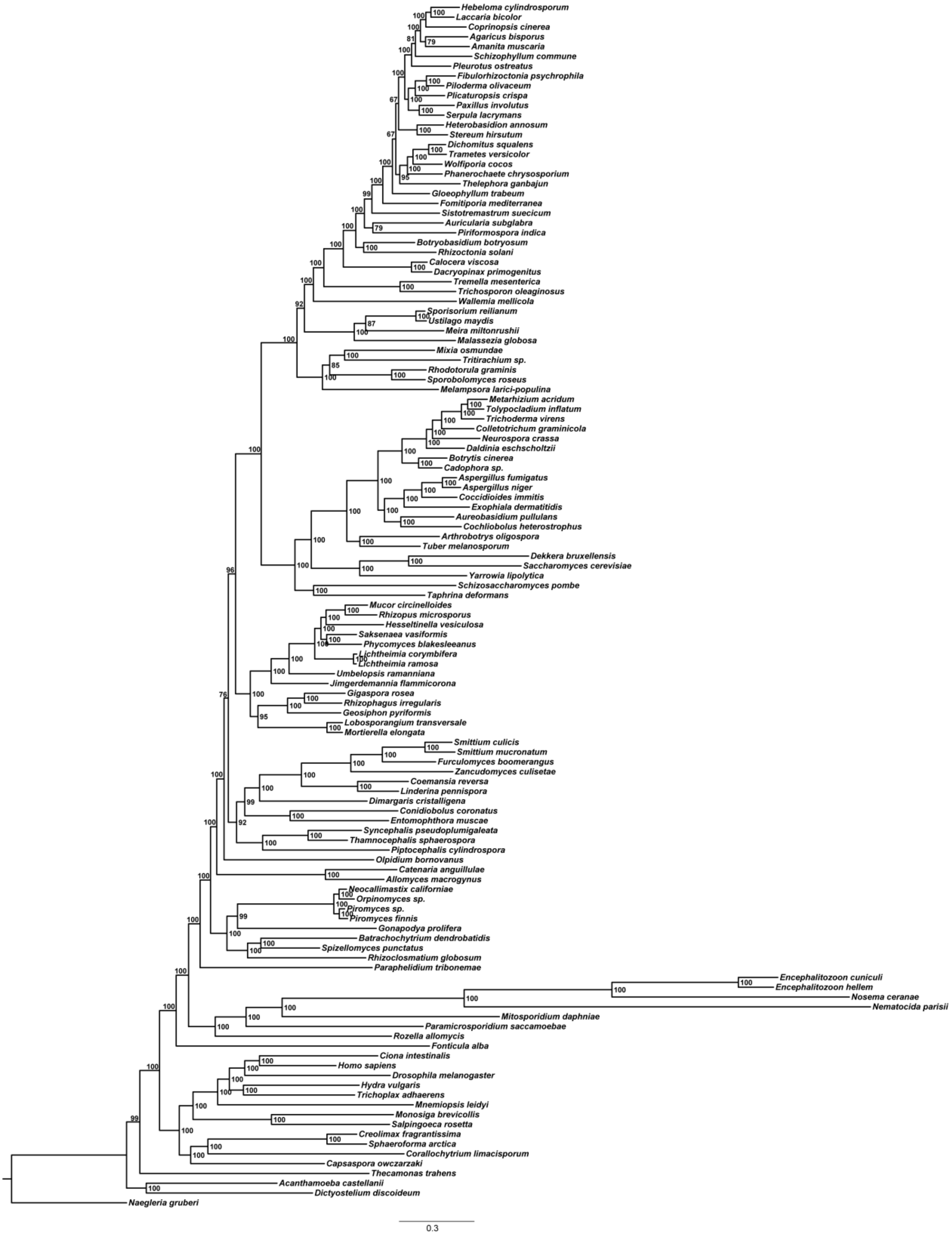


**Supplementary Figure 1. Phylogenetic tree based on 272 single copy orthologous clusters.** The numbers on the branches are ultra-fast bootstrap support values inferred under the LG+G model (ML analysis). The scale bar represents 0.3 expected change per site.

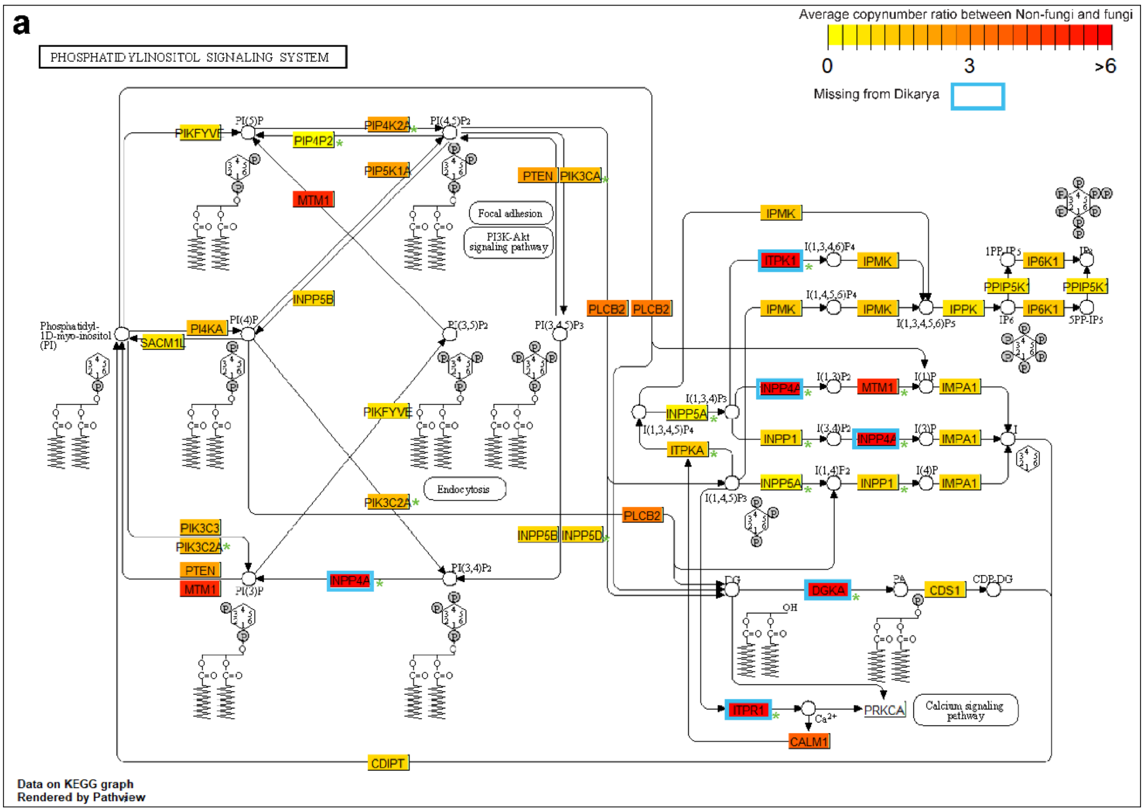


**b**

| GeneID | Gene name | Ratio (P/F) | extinction | name | KEGG Orthology | KEGG ENZYME | Homologus group | Heterolobosea | Amobozoa | Apusozoa | Filasterea | Corallochytrea | Ichtyosporea | Choanoflagellata | Metazoa | Nuclearia | Opisthosporidia | Partr | Chytridiomycota | Blastocladiomycota | Olpid | Zoopagomycota | Mucoromycota | Taphrinomycotina | Saccharomycotina | Pezizomycotina | Pucciniomycotina | Ustilagomycotina | Agaricomycotina |
|---|---|---|---|---|---|---|---|---|---|---|---|---|---|---|---|---|---|---|---|---|---|---|---|---|---|---|---|---|---|
| 22876 | INPP5F | NA | | inositol polyphosphate-5-phosphatase F | KO:K21798 | EC:3.1.3.- | NA | - | - | - | - | - | - | - | - | - | - | - | - | - | - | - | - | - | - | - | - | - | - |
| 5578 | PRKCA | NA | | protein kinase C alpha | KO:K02677 | EC:2.7.11.13 | NA | - | - | - | - | - | - | - | - | - | - | - | - | - | - | - | - | - | - | - | - | - | - |
| 200576 | PIKFYVE | 0.7 | | phosphoinositide kinase FYVE-type zinc finger containing | KO:K00921 | EC:2.7.1.150 | mcl_2006 | 1 | 1 | 1 | 1 | 1 | 0 | 1 | 1 | 1 | 1 | 3 | 2 | 1 | 1 | 1 | 2 | 1 | 1 | 1 | 1 | 1 | 1 |
| 10423 | CDIPT | 1.2 | | CDP-diacylglycerol--inositol 3-phosphatidyltransferase | KO:K00999 | EC:2.7.8.11 | mcl_1478 | 1 | 1 | 2 | 1 | 1 | 2 | 2 | 1 | 1 | 1 | 3 | 1 | 2 | 0 | 1 | 1 | 1 | 1 | 1 | 2 | 1 | 1 |
| 64768 | IPPK | 0.7 | | inositol-pentakisphosphate 2-kinase | KO:K10572 | EC:2.7.1.158 | mcl_3140 | 1 | 1 | 0 | 1 | 1 | 1 | 0 | 1 | 1 | 0 | 0 | 1 | 2 | 1 | 1 | 1 | 1 | 1 | 1 | 1 | 1 | 1 |
| 1040 | CDS1 | 0.9 | | CDP-diacylglycerol synthase 1 | KO:K00981 | EC:2.7.7.41 | mcl_1364 | 0 | 1 | 1 | 1 | 1 | 2 | 1 | 1 | 0 | 1 | 1 | 1 | 3 | 1 | 1 | 2 | 1 | 1 | 1 | 1 | 1 | 1 |
| 3633 | INPP5B | 1.1 | | inositol polyphosphate-5-phosphatase B | KO:K01099 | EC:3.1.3.36 | mcl_199 | 5 | 5 | 5 | 8 | 7 | 5 | 6 | 6 | 5 | 3 | 8 | 7 | 8 | 4 | 4 | 6 | 4 | 4 | 5 | 5 | 5 | 6 |
| 3635 | INPP5D | 1.1 | | inositol polyphosphate-5-phosphatase D | KO:K03084 | EC:3.1.3.86 | mcl_199 | 5 | 5 | 5 | 8 | 7 | 5 | 6 | 6 | 5 | 3 | 8 | 7 | 8 | 4 | 4 | 6 | 4 | 4 | 5 | 5 | 5 | 6 |
| 9677 | PPIP5K1 | 0.7 | | diphosphoinositol pentakisphosphate kinase 1 | KO:K13024 | EC:2.7.4.24 | mcl_1378 | 0 | 0 | 0 | 1 | 1 | 1 | 1 | 2 | 0 | 1 | 3 | 1 | 2 | 1 | 1 | 1 | 1 | 1 | 1 | 1 | 1 | 1 |
| 5290 | PIK3CA | 1.7 | | phosphatidylinositol-4;5-bisphosphate 3-kinase catalytic subunit alpha | KO:K00922 | EC:2.7.1.153 | mcl_492 | 8 | 12 | 5 | 7 | 3 | 2 | 6 | 5 | 2 | 3 | 13 | 3 | 7 | 0 | 3 | 4 | 3 | 3 | 3 | 3 | 3 | 3 |
| 5289 | PIK3C3 | 1.7 | | phosphatidylinositol 3-kinase catalytic subunit type 3 | KO:K00914 | EC:2.7.1.137 | mcl_492 | 8 | 12 | 5 | 7 | 3 | 2 | 6 | 5 | 2 | 3 | 13 | 3 | 7 | 0 | 3 | 4 | 3 | 3 | 3 | 3 | 3 | 3 |
| 5286 | PIK3C2A | 1.7 | | phosphatidylinositol-4-phosphate 3-kinase catalytic subunit type 2 alpha | KO:K00923 | EC:2.7.1.154 | mcl_492 | 8 | 12 | 5 | 7 | 3 | 2 | 6 | 5 | 2 | 3 | 13 | 3 | 7 | 0 | 3 | 4 | 3 | 3 | 3 | 3 | 3 | 3 |
| 5297 | PI4KA | 1.7 | | phosphatidylinositol 4-kinase alpha | KO:K00888 | EC:2.7.1.67 | mcl_492 | 8 | 12 | 5 | 7 | 3 | 2 | 6 | 5 | 2 | 3 | 13 | 3 | 7 | 0 | 3 | 4 | 3 | 3 | 3 | 3 | 3 | 3 |
| 3612 | IMPA1 | 1.2 | | inositol monophosphatase 1 | KO:K01092 | EC:3.1.3.25 | mcl_1090 | 1 | 3 | 5 | 3 | 2 | 3 | 2 | 4 | 2 | 2 | 2 | 2 | 3 | 1 | 3 | 3 | 2 | 2 | 4 | 2 | 2 | 2 |
| 3628 | INPP1 | 1.2 | | inositol polyphosphate-1-phosphatase | KO:K01107 | EC:3.1.3.57 | mcl_1090 | 1 | 3 | 5 | 3 | 2 | 3 | 2 | 4 | 2 | 2 | 2 | 2 | 3 | 1 | 3 | 3 | 2 | 2 | 4 | 2 | 2 | 2 |
| 3706 | ITPKA | 1.3 | | inositol-trisphosphate 3-kinase A | KO:K00911 | EC:2.7.1.127 | mcl_768 | 2 | 4 | 1 | 3 | 2 | 2 | 2 | 4 | 2 | 1 | 4 | 3 | 2 | 1 | 2 | 3 | 2 | 2 | 2 | 2 | 2 | 2 |
| 253430 | IPMK | 1.3 | | inositol polyphosphate multikinase | KO:K00915 | EC:2.7.1.140 2.7.1.151 | mcl_768 | 2 | 4 | 1 | 3 | 2 | 2 | 2 | 4 | 2 | 1 | 4 | 3 | 2 | 1 | 2 | 3 | 2 | 2 | 2 | 2 | 2 | 2 |
| 9807 | IP6K1 | 1.3 | | inositol hexakisphosphate kinase 1 | KO:K07756 | EC:2.7.4.21 | mcl_768 | 2 | 4 | 1 | 3 | 2 | 2 | 2 | 4 | 2 | 1 | 4 | 3 | 2 | 1 | 2 | 3 | 2 | 2 | 2 | 2 | 2 | 2 |
| 5728 | PTEN | 2.3 | | phosphatase and tensin homolog | KO:K01110 | EC:3.1.3.16 3.1.3.48 3.1.3.67 | mcl_1268 | 13 | 14 | 8 | 4 | 5 | 4 | 6 | 9 | 2 | 2 | 12 | 8 | 13 | 3 | 2 | 6 | 2 | 2 | 2 | 2 | 2 | 3 |
| 5305 | PIP4K2A | 2.4 | | phosphatidylinositol-5-phosphate 4-kinase type 2 alpha | KO:K00920 | EC:2.7.1.149 | mcl_586 | 5 | 7 | 8 | 3 | 2 | 2 | 5 | 4 | 1 | 1 | 2 | 4 | 3 | 1 | 1 | 3 | 1 | 1 | 2 | 1 | 1 | 1 |
| 8394 | PIP5K1A | 2.4 | | phosphatidylinositol-4-phosphate 5-kinase type 1 alpha | KO:K00889 | EC:2.7.1.68 | mcl_586 | 5 | 7 | 8 | 3 | 2 | 2 | 5 | 4 | 1 | 1 | 2 | 4 | 3 | 1 | 1 | 3 | 1 | 1 | 2 | 1 | 1 | 1 |
| 5330 | PLCB2 | 3.3 | | phospholipase C beta 2 | KO:K05858 | EC:3.1.4.11 | mcl_214 | 39 | 20 | 19 | 14 | 9 | 12 | 8 | 23 | 1 | 4 | 5 | 4 | 33 | 2 | 6 | 10 | 2 | 2 | 4 | 2 | 3 | 5 |
| 3705 | ITPK1 | 5.9 | 186 | inositol-tetrakisphosphate 1-kinase | KO:K00913 | EC:2.7.1.159 2.7.1.134 | mcl_4632 | 0 | 1 | 1 | 1 | 0 | 0 | 1 | 2 | 1 | 1 | 1 | 1 | 1 | 0 | 1 | 1 | 0 | 0 | 0 | 0 | 0 | 0 |
| 3708 | ITPR1 | 6.0 | 186 | inositol 1;4;5-trisphosphate receptor type 1 | KO:K04958 | NA | mcl_4681 | 5 | 1 | 2 | 2 | 2 | 2 | 3 | 2 | 1 | 0 | 0 | 3 | 0 | 0 | 2 | 1 | 0 | 0 | 0 | 0 | 0 | 0 |
| 801 | CALM1 | 3.8 | | calmodulin 1 | KO:K02183 | NA | mcl_30 | 56 | 35 | 49 | 9 | 14 | 14 | 30 | 61 | 13 | 5 | 28 | 18 | 21 | 7 | 10 | 16 | 8 | 6 | 9 | 8 | 6 | 8 |
| 4534 | MTM1 | 5.0 | | myotubularin 1 | KO:K01108 | EC:3.1.3.64 3.1.3.95 | mcl_694 | 2 | 10 | 3 | 7 | 4 | 3 | 4 | 7 | 3 | 1 | 2 | 1 | 2 | 0 | 1 | 1 | 1 | 1 | 1 | 1 | 1 | 1 |
| 55529 | PIP4P2 | 0.3 | | phosphatidylinositol-4;5-bisphosphate 4-phosphatase 2 | KO:K13084 | EC:3.1.3.78 | mcl_28872 | 0 | 0 | 0 | 0 | 0 | 0 | 0 | 1 | 0 | 0 | 0 | 0 | 0 | 0 | 0 | 0 | 0 | 0 | 0 | 0 | 0 | 0 |
| 3632 | INPP5A | 0.5 | | inositol polyphosphate-5-phosphatase A | KO:K01106 | EC:3.1.3.56 | mcl_15652 | 0 | 0 | 0 | 1 | 0 | 0 | 1 | 1 | 0 | 0 | 0 | 0 | 0 | 0 | 0 | 0 | 0 | 0 | 0 | 0 | 0 | 0 |
| 1606 | DGKA | 6.0 | 186 | diacylglycerol kinase alpha | KO:K00901 | EC:2.7.1.107 | mcl_3323 | 0 | 5 | 2 | 4 | 3 | 3 | 3 | 5 | 1 | 0 | 0 | 3 | 0 | 0 | 2 | 2 | 0 | 0 | 0 | 0 | 0 | 0 |
| 3631 | INPP4A | 6.0 | 159 | inositol polyphosphate-4-phosphatase type I A | KO:K01109 | EC:3.1.3.66 | mcl_7090 | 0 | 0 | 4 | 1 | 0 | 0 | 2 | 2 | 0 | 0 | 0 | 0 | 2 | 0 | 0 | 0 | 0 | 0 | 0 | 0 | 0 | 0 |

**Supplementary Figure 2. Decreased redundancy and diversity of the phosphatidylinositol signalling system in fungi. a)** KEGG graph based on *Homos sapiens* gene IDs. A rectangle can contain multiple genes from the same homologous group, therefore we have only shown the representative gene name that KEGG uses, in the chart and table (b) as well. Colouring of the rectangles is based on the average copy number ratio between non-fungi and Fungi, warmer colour represents more members of a given HG in non-fungi than in Fungi. Green asterisk (*) indicates the absence of a component in the *Saccharomyces* pathway based on the KEGG pathway (sce04070), while blue stroke represents the absence of a component from Dikarya based on our clustering. **b)** The table shows the average copy numbers of the HGs in which the components of the pathway were clustered. The ratio (M/F) was used to colour the rectangles in chart (a).

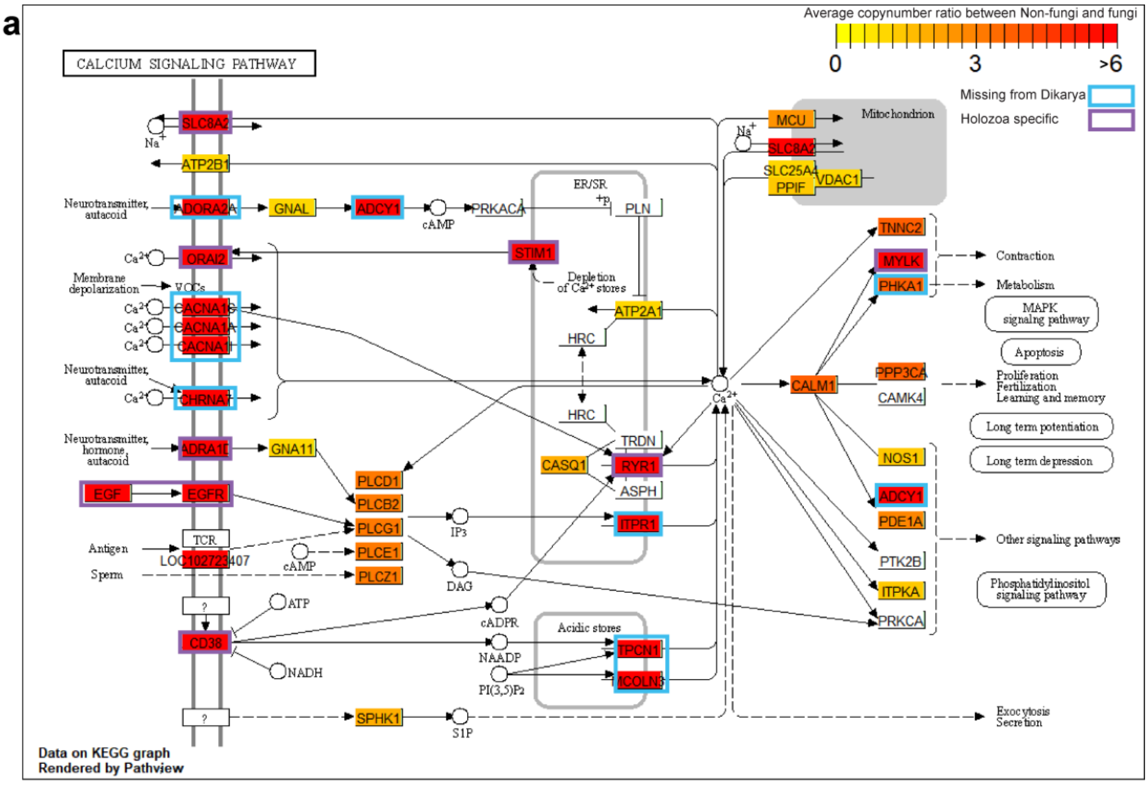


**b**

| GeneID | Gene name | Ratio (P/F) | name | KEGG Orthology | Homologus group | Heterolobosea | Amobozoa | Apusozoa | Filasterea | Corallochytrea | Ichtyosporea | Choanoflagellata | Metazoa | Nuclearia | Opisthosporidia | Partr | Chytridiomycota | Blastocladiomycota | Olpid | Zoopagomycota | Mucoromycota | Taphrinomycotina | Saccharomycotina | Pezizomycotina | Pucciniomycotina | Ustilagomycotina | Agaricomycotina |
|---|---|---|---|---|---|---|---|---|---|---|---|---|---|---|---|---|---|---|---|---|---|---|---|---|---|---|---|
| 5255 | PHKA1 | 3.88 | PHKA1; phosphorylase kinase regulatory subunit alpha 1 | KO:K07190 | mcl_3995 | 0 | 1 | 0 | 2 | 2 | 1.5 | 0 | 2.17 | 0 | 1 | 5 | 1 | 2 | 1 | 1 | 1.5 | 0 | 0 | 0 | 0 | 0 | 0 |
| 135 | ADORA2A | 6 | ADORA2A; adenosine A2a receptor | KO:K04266 | mcl_183 | 0 | 0 | 0 | 0 | 0 | 0 | 0 | 140 | 0 | 1 | 0 | 7.5 | 0 | 0 | 0 | 0 | 0 | 0 | 0 | 0 | 0 | 0 |
| 107 | ADCY1 | 6 | ADCY1; adenylate cyclase 1 | KO:K08041 | mcl_4360 | 0 | 1 | 1 | 0 | 0 | 1 | 0 | 5.83 | 0 | 1 | 7 | 1 | 2 | 0 | 0 | 0 | 0 | 0 | 0 | 0 | 0 | 0 |
| 775 | CACNA1C | 6 | CACNA1C; calcium voltage-gated channel subunit alpha1 C | KO:K04850 | mcl_3251 | 0 | 0 | 5 | 0 | 2 | 0 | 2 | 12.2 | 0 | 0 | 0 | 1.33 | 0 | 0 | 0 | 0 | 0 | 0 | 0 | 0 | 0 | 0 |
| 773 | CACNA1A | 6 | CACNA1A; calcium voltage-gated channel subunit alpha1 A | KO:K04344 | mcl_3251 | 0 | 0 | 5 | 0 | 2 | 0 | 2 | 12.2 | 0 | 0 | 0 | 1.33 | 0 | 0 | 0 | 0 | 0 | 0 | 0 | 0 | 0 | 0 |
| 8911 | CACNA1I | 6 | CACNA1I; calcium voltage-gated channel subunit alpha1 I | KO:K04856 | mcl_3251 | 0 | 0 | 5 | 0 | 2 | 0 | 2 | 12.2 | 0 | 0 | 0 | 1.33 | 0 | 0 | 0 | 0 | 0 | 0 | 0 | 0 | 0 | 0 |
| 1139 | CHRNA7 | 6 | CHRNA7; cholinergic receptor nicotinic alpha 7 subunit | KO:K04809 | mcl_2375 | 0 | 0 | 0 | 1 | 0 | 2 | 3 | 29.5 | 0 | 0 | 2 | 0 | 0 | 0 | 0 | 0 | 0 | 0 | 0 | 0 | 0 | 1 |
| 146 | ADRA1D | 6 | ADRA1D; adrenoceptor alpha 1D | KO:K04137 | mcl_72 | 0 | 0 | 0 | 0 | 0 | 0 | 0 | 188 | 0 | 0 | 0 | 3 | 0 | 0 | 0 | 0 | 0 | 0 | 0 | 0 | 0 | 0 |
| 3708 | ITPR1 | 6 | ITPR1; inositol 1;4;5-trisphosphate receptor type 1 | KO:K04958 | mcl_4681 | 5 | 1 | 2 | 2 | 2 | 2 | 3 | 1.8 | 1 | 0 | 0 | 3 | 0 | 0 | 1.5 | 1 | 0 | 0 | 0 | 0 | 0 | 0 |
| 53373 | TPCN1 | 6 | TPCN1; two pore segment channel 1 | KO:K16896 | mcl_5883 | 0 | 1.5 | 6 | 1 | 3 | 2.5 | 5.5 | 2.8 | 0 | 0 | 0 | 0 | 0 | 0 | 0 | 0 | 0 | 0 | 0 | 0 | 0 | 0 |
| 55283 | MCOLN3 | 6 | MCOLN3; mucolipin TRP cation channel 3 | KO:K04994 | mcl_10410 | 2 | 1 | 0 | 1 | 0 | 0 | 1 | 1.83 | 0 | 0 | 0 | 0 | 0 | 0 | 0 | 0 | 0 | 0 | 0 | 0 | 0 | 0 |
| 6543 | SLC8A2 | 6 | SLC8A2; solute carrier family 8 member A2 | KO:K05849 | mcl_7999 | 0 | 0 | 0 | 0 | 0 | 0 | 2 | 2.33 | 0 | 0 | 0 | 0 | 0 | 0 | 0 | 0 | 0 | 0 | 0 | 0 | 0 | 0 |
| 80228 | ORAI2 | 6 | ORAI2; ORAI calcium release-activated calcium modulator 2 | KO:K16057 | mcl_10858 | 0 | 0 | 0 | 1 | 0 | 2 | 0 | 1.6 | 0 | 0 | 0 | 0 | 0 | 0 | 0 | 0 | 0 | 0 | 0 | 0 | 0 | 0 |
| 6786 | STIM1 | 6 | STIM1; stromal interaction molecule 1 | KO:K16059 | mcl_12541 | 0 | 0 | 0 | 1 | 0 | 1 | 1 | 1.17 | 0 | 0 | 0 | 0 | 0 | 0 | 0 | 0 | 0 | 0 | 0 | 0 | 0 | 0 |
| 1950 | EGF | 6 | EGF; epidermal growth factor | KO:K04357 | mcl_5247 | 0 | 0 | 0 | 0 | 0 | 0 | 0 | 15.3 | 0 | 0 | 0 | 0 | 0 | 0 | 0 | 0 | 0 | 0 | 0 | 0 | 0 | 0 |
| 1956 | EGFR | 6 | EGFR; epidermal growth factor receptor | KO:K04361 | mcl_9121 | 0 | 0 | 0 | 0 | 1 | 7.5 | 3.5 | 4.33 | 0 | 0 | 0 | 0 | 0 | 0 | 0 | 0 | 0 | 0 | 0 | 0 | 0 | 0 |
| 952 | CD38 | 6 | CD38; CD38 molecule | KO:K01242 | mcl_11147 | 0 | 0 | 0 | 1 | 0 | 0 | 0 | 4 | 0 | 0 | 0 | 0 | 0 | 0 | 0 | 0 | 0 | 0 | 0 | 0 | 0 | 0 |
| 6261 | RYR1 | 6 | RYR1; ryanodine receptor 1 | KO:K04961 | mcl_17274 | 0 | 0 | 0 | 1 | 0 | 0 | 0 | 1.5 | 0 | 0 | 0 | 0 | 0 | 0 | 0 | 0 | 0 | 0 | 0 | 0 | 0 | 0 |
| 4638 | MYLK | 6 | MYLK; myosin light chain kinase | KO:K00907 | mcl_25532 | 0 | 0 | 0 | 0 | 0 | 0 | 0 | 2 | 0 | 0 | 0 | 0 | 0 | 0 | 0 | 0 | 0 | 0 | 0 | 0 | 0 | 0 |
| 490 | ATP2B1 | 0.94 | ATP2B1; ATPase plasma membrane Ca2+ transporting 1 | KO:K05850 | mcl_80 | 11 | 5.5 | 9 | 9 | 5 | 5.5 | 8 | 8.67 | 3 | 5.67 | 17 | 11.1 | 18 | 10 | 8.83 | 12.9 | 4 | 4 | 9.31 | 5.4 | 6.25 | 5.32 |
| 487 | ATP2A1 | 0.94 | ATP2A1; ATPase sarcoplasmic/endoplasmic reticulum Ca2+ transpo | KO:K05853 | mcl_80 | 11 | 5.5 | 9 | 9 | 5 | 5.5 | 8 | 8.67 | 3 | 5.67 | 17 | 11.1 | 18 | 10 | 8.83 | 12.9 | 4 | 4 | 9.31 | 5.4 | 6.25 | 5.32 |
| 2774 | GNAL | 1.04 | GNAL; G protein subunit alpha L | KO:K04633 | mcl_84 | 28 | 10.5 | 0 | 8 | 3 | 6 | 2.5 | 9.17 | 4 | 3 | 12 | 11.6 | 14 | 5 | 2.92 | 8.71 | 2.5 | 2 | 3.13 | 5.2 | 3.25 | 13.4 |
| 2767 | GNA11 | 1.04 | GNA11; G protein subunit alpha 11 | KO:K04635 | mcl_84 | 28 | 10.5 | 0 | 8 | 3 | 6 | 2.5 | 9.17 | 4 | 3 | 12 | 11.6 | 14 | 5 | 2.92 | 8.71 | 2.5 | 2 | 3.13 | 5.2 | 3.25 | 13.4 |
| 291 | SLC25A4 | 1.09 | SLC25A4; solute carrier family 25 member 4 | KO:K05863 | mcl_10 | 45 | 45.5 | 38 | 38 | 35 | 38.5 | 34.5 | 36.5 | 23 | 20 | 62 | 29.1 | 56.5 | 16 | 32.9 | 47.1 | 26 | 33 | 37.3 | 31.2 | 31.3 | 34.4 |
| 7416 | VDAC1 | 1.09 | VDAC1; voltage dependent anion channel 1 | KO:K05862 | mcl_1136 | 1 | 1 | 1 | 1 | 1 | 1 | 1 | 1.83 | 1 | 1 | 1 | 1.38 | 2.5 | 1 | 1.33 | 2 | 1 | 1.33 | 1 | 1 | 1 | 1.06 |
| 5530 | PPP3CA | 1.12 | PPP3CA; protein phosphatase 3 catalytic subunit alpha | KO:K04348 | mcl_67 | 9 | 9.5 | 10 | 7 | 8 | 10 | 12 | 10.8 | 9 | 4.43 | 14 | 10.3 | 15 | 5 | 7.92 | 11.7 | 8 | 11 | 8.38 | 7.6 | 7.75 | 8.97 |
| 10105 | PPIF | 1.25 | PPIF; peptidylprolyl isomerase F | KO:K09565 | mcl_47 | 19 | 13 | 17 | 11 | 6 | 15 | 15 | 17.5 | 11 | 4.29 | 21 | 15.6 | 21 | 7 | 12.7 | 11.9 | 9 | 9 | 10.5 | 12 | 9.5 | 13.5 |
| 4842 | NOS1 | 1.34 | NOS1; nitric oxide synthase 1 | KO:K13240 | mcl_209 | 7 | 6.5 | 5 | 3 | 4 | 6.5 | 4 | 5.67 | 4 | 2.29 | 2 | 3.88 | 5.5 | 6 | 4.42 | 5.21 | 4 | 3 | 5 | 2.8 | 3 | 3.52 |
| 3706 | ITPKA | 1.34 | ITPKA; inositol-trisphosphate 3-kinase A | KO:K00911 | mcl_768 | 2 | 4 | 1 | 3 | 2 | 2 | 2 | 3.67 | 2 | 1 | 4 | 2.75 | 2 | 1 | 2.18 | 2.93 | 2 | 2 | 1.88 | 1.8 | 2 | 2.23 |
| 844 | CASQ1 | 1.84 | CASQ1; calsequestrin 1 | KO:K23468 | mcl_576 | 2 | 3.5 | 4 | 2 | 1 | 4 | 6 | 6.83 | 1 | 1.5 | 7 | 5 | 4 | 1 | 2.42 | 2.57 | 2.5 | 1.67 | 2.5 | 2.4 | 1.75 | 2.19 |
| 8877 | SPHK1 | 1.96 | SPHK1; sphingosine kinase 1 | KO:K04718 | mcl_651 | 4 | 4 | 4 | 4 | 4 | 5 | 2.5 | 3.17 | 0 | 1 | 1 | 1.5 | 2 | 1 | 1.78 | 2.21 | 1 | 2 | 2.13 | 2 | 2 | 2 |
| 90550 | MCU | 2.5 | MCU; mitochondrial calcium uniporter | KO:K20858 | mcl_5782 | 1 | 1 | 2 | 1 | 1 | 1 | 1 | 1.2 | 1 | 0 | 1 | 1.5 | 2 | 0 | 0 | 0 | 1 | 0 | 1 | 0 | 0 | 1 |
| 5136 | PDE1A | 2.93 | PDE1A; phosphodiesterase 1A | KO:K13755 | mcl_271 | 7 | 2.5 | 11 | 3 | 6 | 2 | 6 | 11.5 | 2 | 3 | 8 | 8.13 | 14 | 6 | 1.64 | 4.79 | 1 | 1 | 1.06 | 1 | 0 | 1 |
| 5333 | PLCD1 | 3.29 | PLCD1; phospholipase C delta 1 | KO:K05857 | **mcl_214** | 39 | 19.5 | 19 | 14 | 9 | 11.5 | 8 | 23.3 | 1 | 4 | 5 | 3.71 | 32.5 | 2 | 6.18 | 10.4 | 2 | 2 | 4 | 2.25 | 2.75 | 5.13 |
| 5330 | PLCB2 | 3.29 | PLCB2; phospholipase C beta 2 | KO:K05858 | **mcl_214** | 39 | 19.5 | 19 | 14 | 9 | 11.5 | 8 | 23.3 | 1 | 4 | 5 | 3.71 | 32.5 | 2 | 6.18 | 10.4 | 2 | 2 | 4 | 2.25 | 2.75 | 5.13 |
| 5335 | PLCG1 | 3.29 | PLCG1; phospholipase C gamma 1 | KO:K01116 | **mcl_214** | 39 | 19.5 | 19 | 14 | 9 | 11.5 | 8 | 23.3 | 1 | 4 | 5 | 3.71 | 32.5 | 2 | 6.18 | 10.4 | 2 | 2 | 4 | 2.25 | 2.75 | 5.13 |
| 51196 | PLCE1 | 3.29 | PLCE1; phospholipase C epsilon 1 | KO:K05860 | **mcl_214** | 39 | 19.5 | 19 | 14 | 9 | 11.5 | 8 | 23.3 | 1 | 4 | 5 | 3.71 | 32.5 | 2 | 6.18 | 10.4 | 2 | 2 | 4 | 2.25 | 2.75 | 5.13 |
| 89869 | PLCZ1 | 3.29 | PLCZ1; phospholipase C zeta 1 | KO:K05861 | **mcl_214** | 39 | 19.5 | 19 | 14 | 9 | 11.5 | 8 | 23.3 | 1 | 4 | 5 | 3.71 | 32.5 | 2 | 6.18 | 10.4 | 2 | 2 | 4 | 2.25 | 2.75 | 5.13 |
| 801 | CALM1 | 3.75 | CALM1; calmodulin 1 | KO:K02183 | mcl_30 | 56 | 35 | 49 | 9 | 14 | 13.5 | 29.5 | 61.3 | 13 | 4.57 | 28 | 18.4 | 20.5 | 7 | 10.3 | 15.9 | 7.5 | 6.33 | 9.19 | 7.6 | 6.25 | 8.32 |
| 7125 | TNNC2 | 3.75 | TNNC2; troponin C2; fast skeletal type | KO:K12042 | mcl_30 | 56 | 35 | 49 | 9 | 14 | 13.5 | 29.5 | 61.3 | 13 | 4.57 | 28 | 18.4 | 20.5 | 7 | 10.3 | 15.9 | 7.5 | 6.33 | 9.19 | 7.6 | 6.25 | 8.32 |

**Supplementary Figure 3. Decreased redundancy and diversity of the calcium signalling system in fungi. a)** KEGG graph based on *Homos sapiens* gene IDs. A rectangle can contain multiple genes from the same homologous group, therefore we have only shown the representative gene name that KEGG uses, in the chart and table (b) as well. Colouring of the rectangles is based on the average copy number ratio between non-fungi and Fungi, warmer colour represents more members of a given HG in non-fungi than in Fungi. Blue stroke represents the absence of a component from Dikarya while purple stroke represents Holozoa specific genes based on our clustering. **b)** The table shows the average copy numbers of the HGs in which the components of the pathway were clustered. The ratio (P/F) was used to colour the rectangles in chart (a).

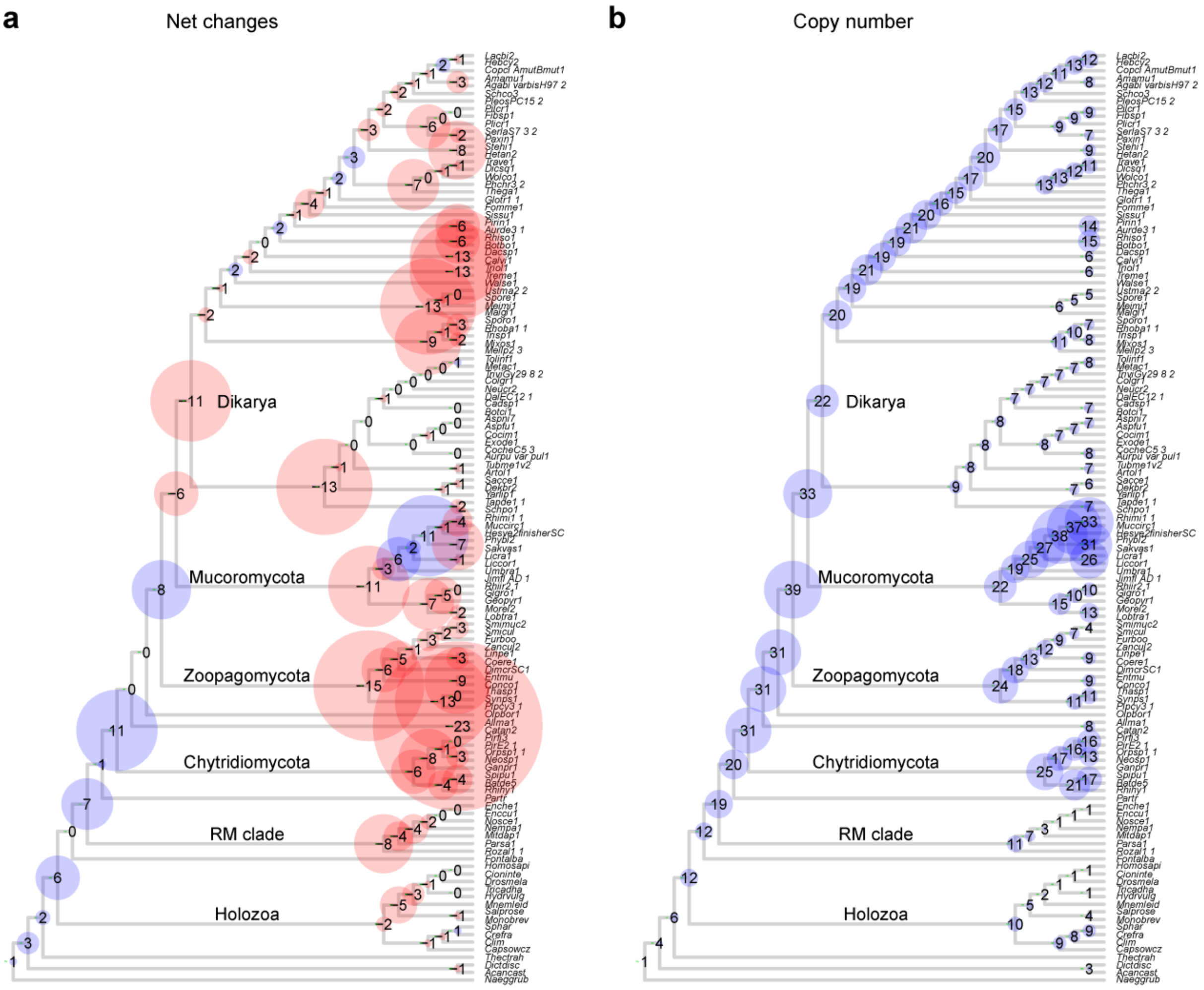


**Supplementary Figure 4. Copy numbers and evolution of the PCL cyclin family. a)** Net changes (expansion - blue; contraction - red) to ancestral protein coding capacity across the fungal phylogeny as inferred by Dollo mapping of duplications and losses. **b)** inferred ancestral copy numbers. Duplications mapping to terminals (that is, resulting in species specific paralogs) are not shown. The size of the circles is proportional to the number of net gain events and copy numbers.

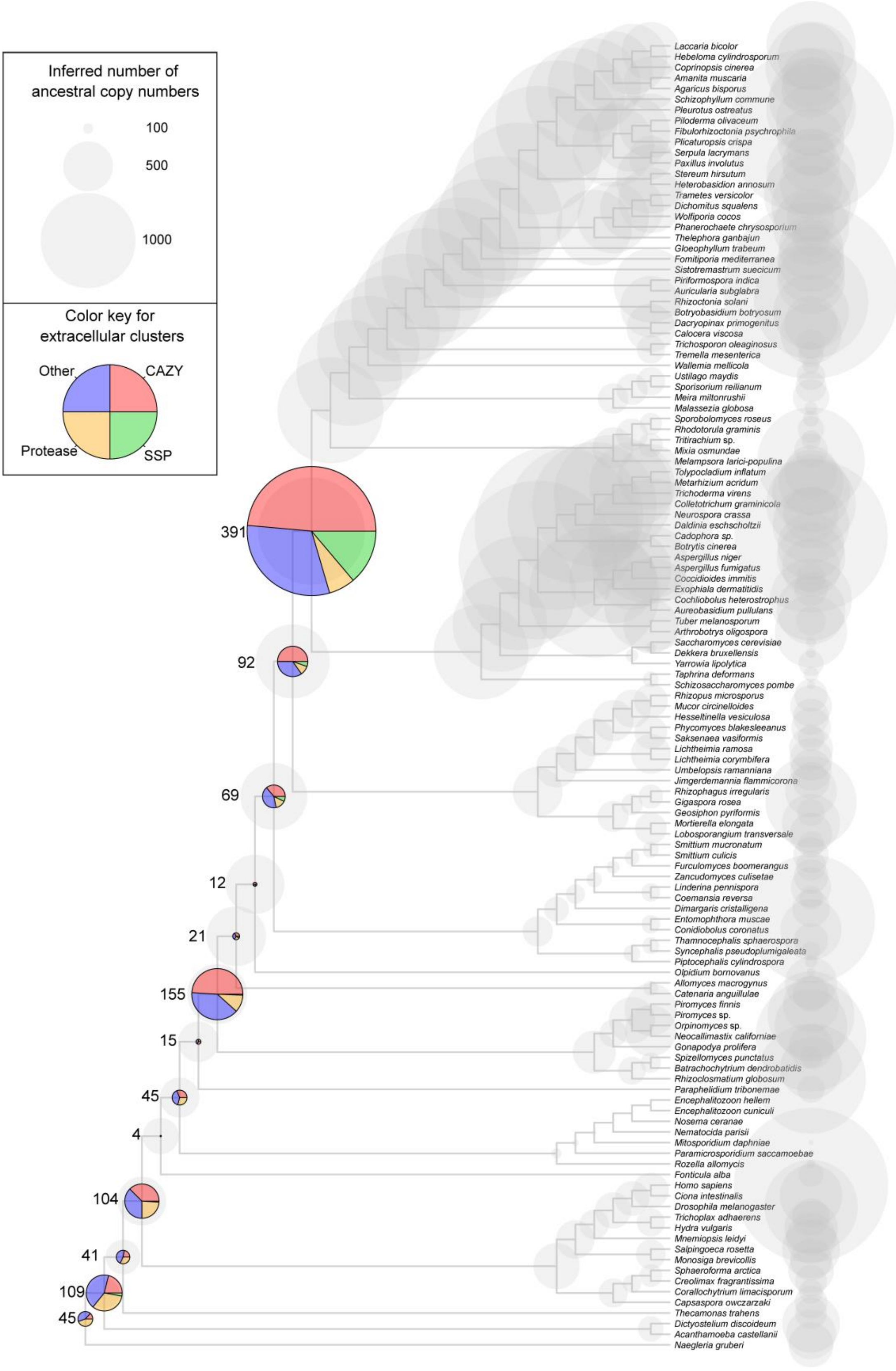


**Supplementary Figure 5. Copy numbers and evolution of homologous groups containing extracellular proteins.** Copy numbers are shown as grey circles, while numbers of duplications are represented by pie-charts. Numbers next to pie charts means the number of duplications. The size of the circles is proportional to the number of net gain events and copy numbers.

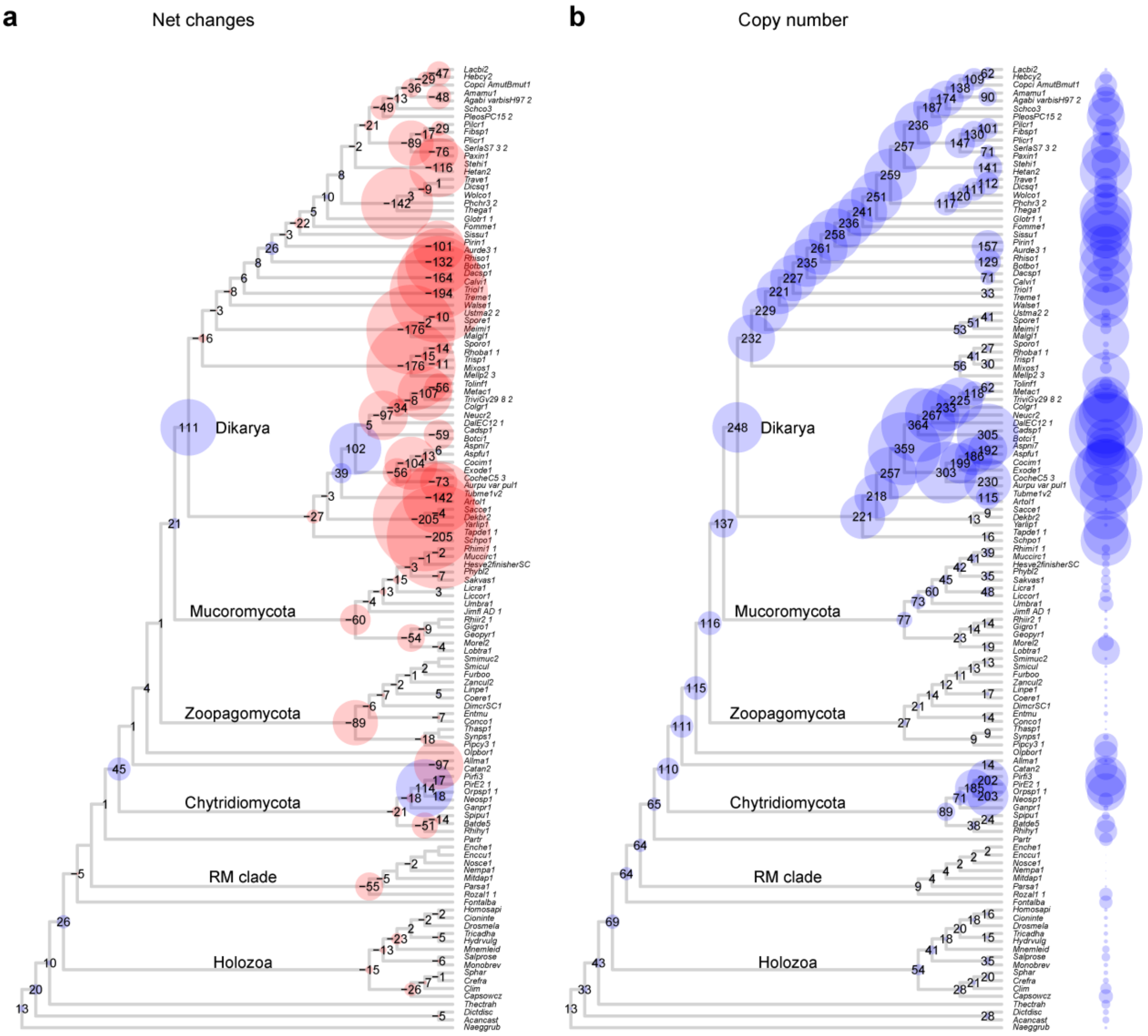


**Supplementary Figure 6. Ancestral copy numbers and evolution of plant cell wall degrading enzyme (PCWDE) HGs. a)** Net changes (expansion - blue; contraction - red) and **b)** inferred ancestral protein coding capacity across the fungal phylogeny. Duplications mapping to terminals (i.e. species specific paralogs) are not shown. The size of the circles is proportional to the number of net gain events and copy numbers in panel (a) and (b), respectively.

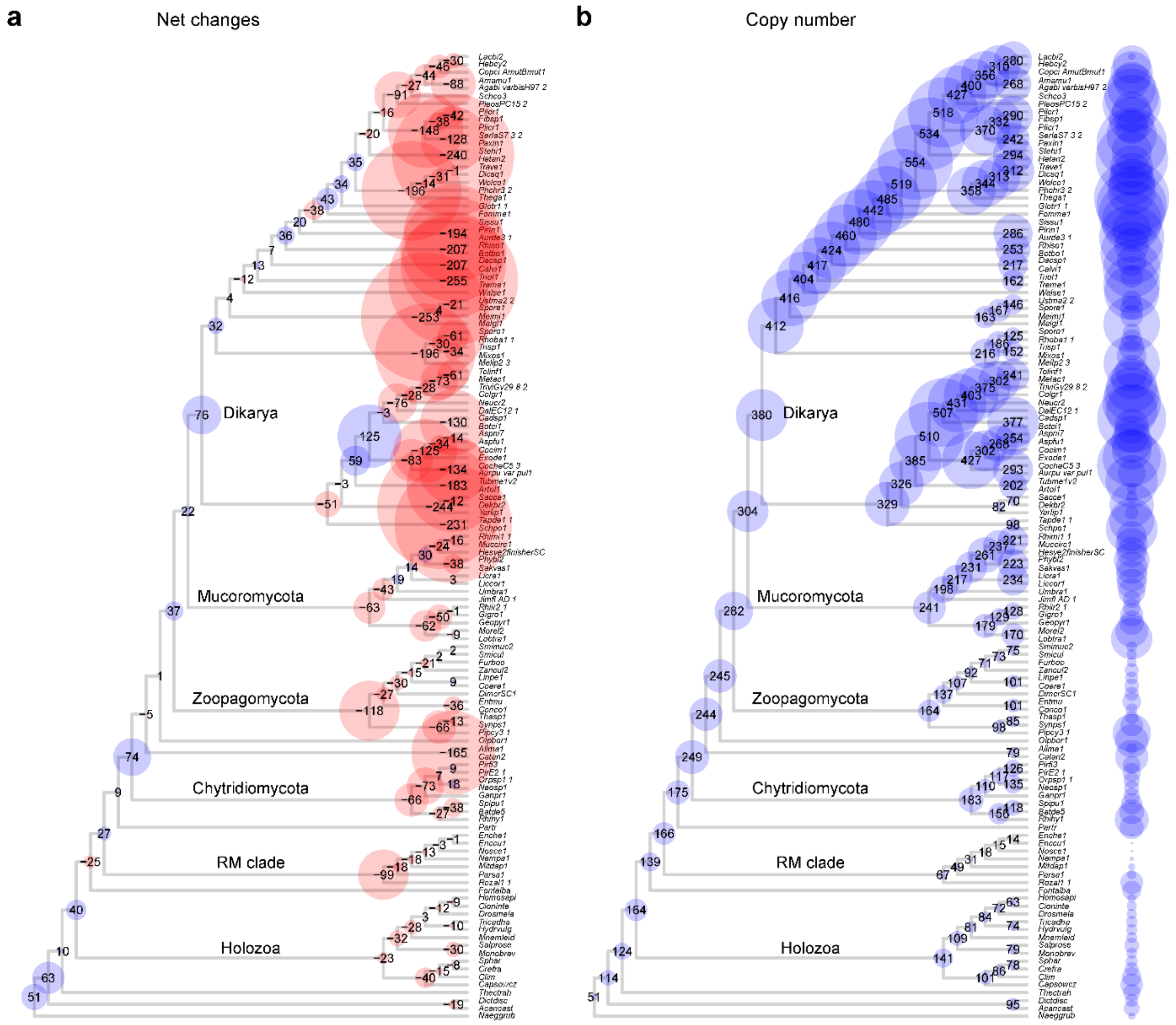


**Supplementary Figure 7. Ancestral copy numbers and evolution of fungal cell wall (FCW) and related cell surface proteins. a)** Net changes (expansion - blue; contraction - red) and **b)** inferred ancestral protein coding capacity across the fungal phylogeny. Duplications mapping to terminals (i.e. species specific paralogs) are not shown. The size of the circles is proportional to the number of net gain events and copy numbers in panel (a) and (b), respectively.

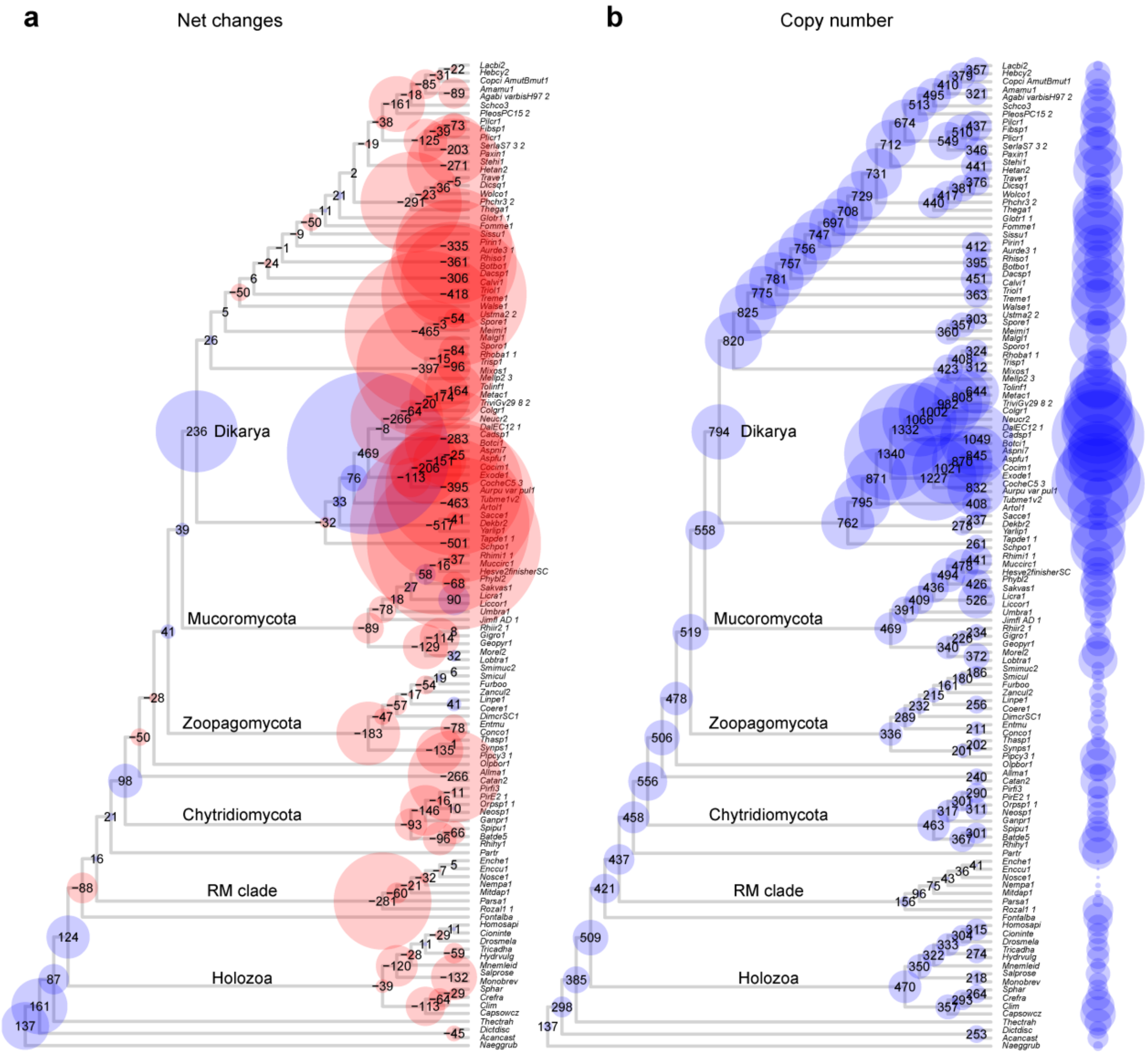


**Supplementary Figure 8. Ancestral copy numbers and evolution of transporters. a)** Net changes (expansion - blue; contraction - red) to ancestral protein coding capacity across the fungal phylogeny. **b)** inferred ancestral copy numbers. Duplications mapping to terminals (i.e. species specific paralogs) are not shown. The size of the circles is proportional to the number of net gain events and copy numbers in panel (a) and (b), respectively.

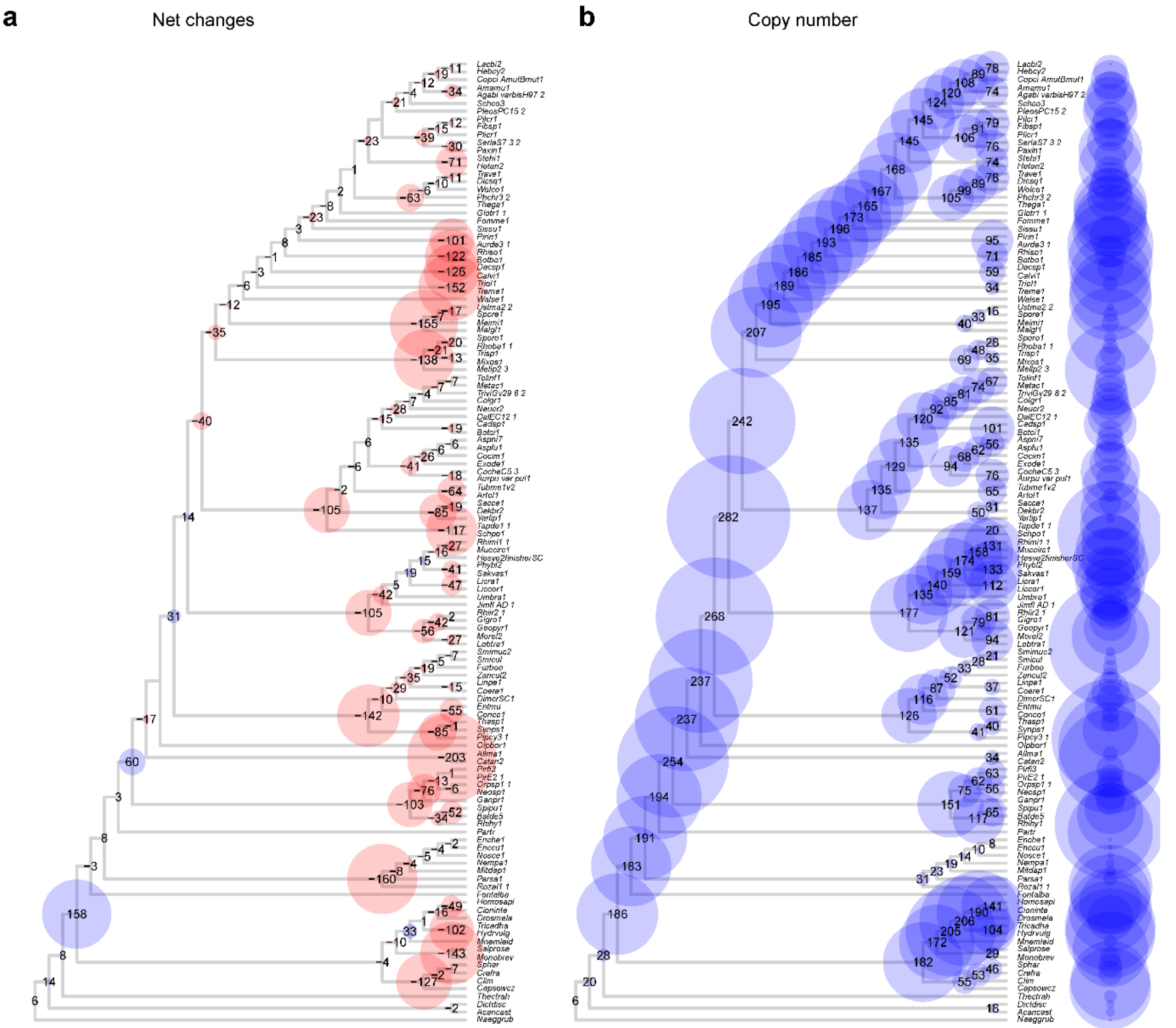


**Supplementary Figure 9. Ancestral copy numbers and evolution of the C2H2 transcription factor family. a)** Net changes (expansion - blue; contraction - red) to ancestral protein coding capacity across the fungal phylogeny. **b)** inferred ancestral copy numbers. Duplications mapping to terminals (i.e. species specific paralogs) are not shown. The size of the circles is proportional to the number of net gain events and copy numbers in panel (a) and (b), respectively.

### Legends for supplementary data

**Supplemental Data 1.** Taxon list of 123 species used for our analyses. BUSCO percentages were obtained with the database 'eukaryote'.

**Supplementary Data 2a.** Conserved HGs shared between protists and early diverging fungi (EDF). 540 HGs which showed at least 70% conservation in any of the groups defined in Supplementary Data 1, emerged before node 140 (split of Nuclearia) and were lost along the backbone between nodes 141 (split of RM clade) and 186 (MRCA of Dikarya).

**Supplemental Data 2b.** Enriched functional categories among HGs (540) lost during fungal evolution. The matrix contains only the significantly enriched (p-value <= 0.05) terms for each backbone node of the tree. Abbreviation n.s. means not significant.

**Supplementary Data 3.** Flagellum-associated proteins. We used the list of flagellar proteins of *Chlamydomonas reinhardtii* from (http://jcb.rupress.org/content/170/1/103.full) as queries in MMSeqs reciprocal best hit searches in the proteomes of the examined species. Numbers in a matrix are -log10 transformed e-values. Species highlighted with green possess flagella.

**Supplementary Data 4a.** Novel core HGs in fungi. 163 HGs which showed at least 70% conservation and were gained along the fungal backbone (>= node 141 =< node 186).

**Supplementary Data 4b.** Homologous groups containing fungal-specific domain, and emerged in the backbone (>= node 141 =< node 186).

**Supplementary Data 5a.** TopGO functional enrichment for the emergence of orthogroups (duplication or de novo birth). Values in the matrix represent the p-value of Fisher's exact test with weight01Fisher algorithm of topGO. Only values < 0.05 are shown. 'n.s.' means not significant.

**Supplementary Data 5b.** TopGO functional enrichment for the 540 orthogroups shared between protists and EDF. The node and clade names represent the last ancestor in which these orthogroups are predicted to be still present. Values in the matrix represent the p-value of Fisher's exact test with weight01Fisher algorithm of topGO. Only values < 0.05 were visualised. 'n.s.' means not significant.

**Supplementary Data 6a.** The dynamics of expansion/contraction of HGs during early fungal evolution. To assess the dynamics of HG changes along the fungal backbone, net change relative to the copy number of the preceding node was calculated and summed across eight nodes from LUFA to the MRCA of Dikarya. We ordered our table based on this score ('Sum of relative net change').

**Supplementary Data 6b.** Inferred gain, loss events and copy numbers of all HGs in the MRCA of Chytridiomycota and other fungi. 27.2% of gains are due to the gene families analysed.

**Supplementary Data 6c.** Inferred gain, loss events and copy numbers of all HGs in the MRCA of Dikarya. 40.7% of gains are attributable to the analysed gene families.

**Supplementary Data 7.** 253 identified transporter HGs, their classification and distribution among clades.

**Supplemental Data 8a.** Evolutionary dynamics of transcription factor families. Column "cumulative net change" means the sum of normalised net changes (gains minus losses) by the copy number of the parental node. The first 12 families (in bold) were visualised in Fig. 4.

**Supplementary Data 8b.** 657 identified TF HGs, their classification and distribution among clades.
